# FAK activity sustains intrinsic and acquired ovarian cancer resistance to platinum chemotherapy

**DOI:** 10.1101/594184

**Authors:** Carlos J. Díaz Osterman, Duygu Ozmadenci, Elizabeth G. Kleinschmidt, Kristin N. Taylor, Allison M. Barrie, Shulin Jiang, Lisa M. Bean, Florian J. Sulzmaier, Christine Jean, Isabelle Tancioni, Kristen Anderson, Sean Uryu, Edward A. Cordasco, Jian Li, Xiao Lei Chen, Guo Fu, Marjaana Ojalill, Pekka Rappu, Jyrki Heino, Adam M. Mark, Guorong Xu, Kathleen M. Fisch, Vihren N. Kolev, David T. Weaver, Jonathan A. Pachter, Balázs Győrffy, Michael T. McHale, Denise C. Connolly, Alfredo Molinolo, Dwayne G. Stupack, David D. Schlaepfer

**Author notes:** authors contributed equally to this study (Carlos J. Díaz Osterman, Duygu Ozmadenci, Elizabeth G. Kleinschmidt, Kristin N. Taylor, Allison M. Barrie, Shulin Jiang, Lisa M. Bean, Florian J. Sulzmaier). Co-corresponding authors: David D. Schlaepfer, Ph.D.; Dwayne G. Stupack, Ph.D., Department of Obstetrics, Gynecology, and Reproductive Sciences UCSD Moores Cancer Center 3855 Health Sciences Dr., MC 0803 La Jolla, CA 92093. INSERM UMR1037, Centre de Recherches en Cancérologie de Toulouse, France (Christine Jean).

## Abstract

Gene copy number changes, cancer stem cell (CSC) increases, and platinum chemotherapy resistance contribute to poor prognosis in patients with recurrent high grade serous ovarian cancer (HGSOC). CSC phenotypes involving Wnt-β-catenin and aldehyde dehydrogenase activities, platinum resistance, and tumor initiating frequency are here associated with spontaneous genetic gains, including genes encoding <u>*K*</u>*RAS*, <u>*M*</u>*YC* and <u>*F*</u>*AK*, in a new murine model of ovarian cancer (KMF). Noncanonical FAK signaling was sufficient to sustain human and KMF tumorsphere proliferation, CSC survival, and platinum resistance. Increased FAK tyrosine phosphorylation occurred in HGSOC patient tumors surviving neo-adjuvant platinum and paclitaxel chemotherapy and platinum resistant tumorspheres acquired FAK dependence for growth. Importantly, combining a pharmacologic FAK inhibitor with platinum overcame chemoresistance and triggered apoptosis *in vitro* and *in vivo*. Knockout, rescue, genomic and transcriptomic analyses collectively identified more than 400 genes regulated along a FAK/β-catenin/Myc axis impacting stemness and DNA repair in HGSOC, with 66 genes gained in a majority of Cancer Genome Atlas samples. Together, these results support combinatorial testing of FAK inhibitors for the treatment of recurrent ovarian cancer.

**Graphical Summary:** 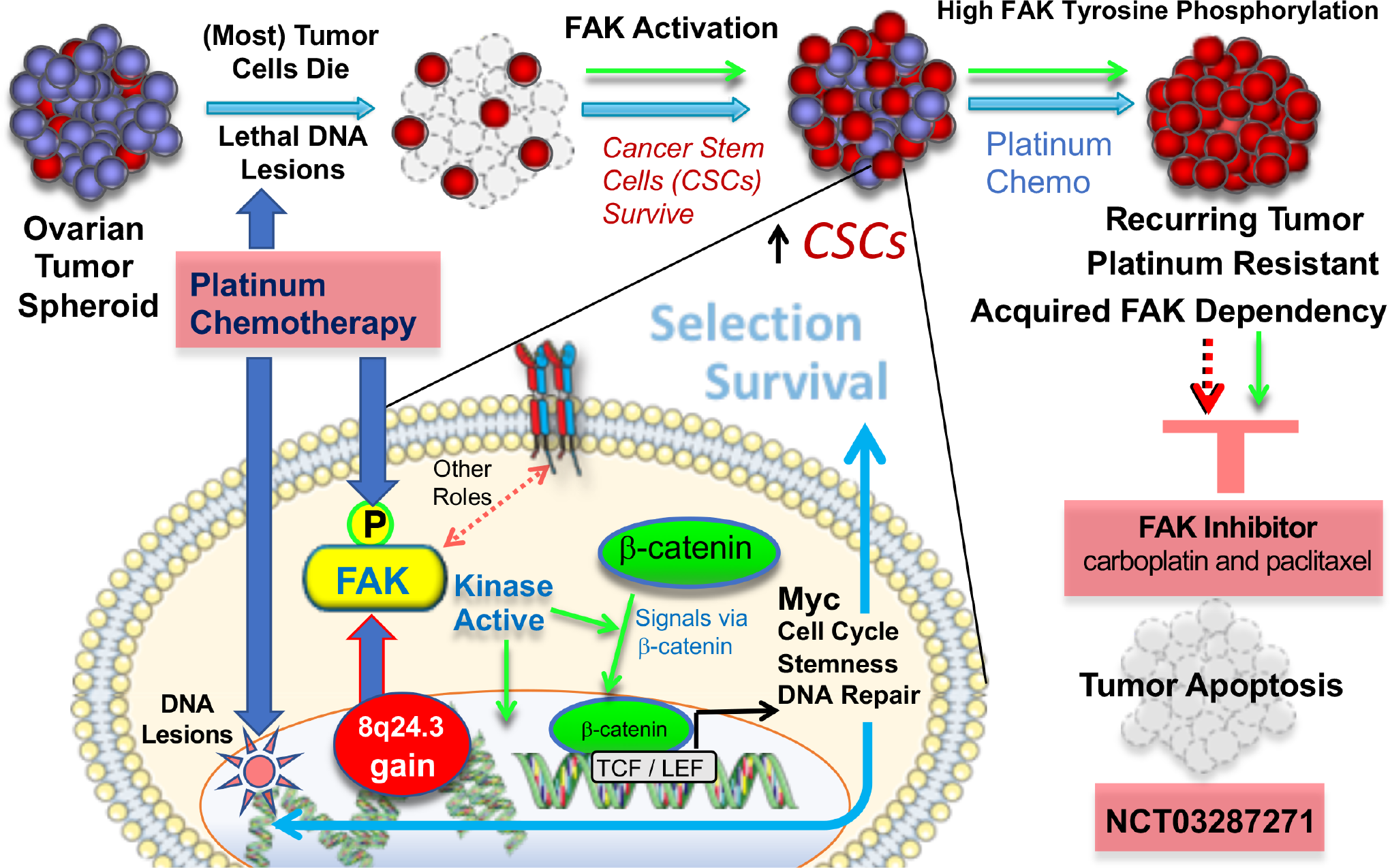

**Key Points:**

- High grade serous ovarian carcinoma tumors contain *PTK2* (FAK) 8q24.3 gains associated with prognostic differences.
- KMF, a new murine ovarian cancer model with <u>K</u>-Ras, <u>M</u>yc, and <u>F</u>AK gene gains and intrinsic platinum resistance.
- FAK activation in tumors surviving platinum chemotherapy promotes cancer stem cell survival.
- FAK facilitates a β-catenin-Myc signaling axis controlling gene expression supporting platinum resistance.
- FAK activity is essential for KMF tumor growth and is a targetable cellular adaptation of platinum resistance.

## Introduction

Ovarian carcinoma is the most lethal gynecologic malignancy in the United States (Siegel, Miller et al., 2018). High grade serous ovarian carcinoma (HGSOC), the most common histologic tumor subtype (Matulonis, Sood et al., 2016), is primarily treated with cytoreductive surgery followed by carboplatin and paclitaxel chemotherapy. The probability of cure is highly dependent on elimination of microscopic disease (Narod, 2016). Approximately 80% of patients with HGSOC will exhibit serial disease recurrence, develop resistance to platinum chemotherapy, and die (Bowtell, Bohm et al., 2015). Although platinum chemotherapy is effective at creating DNA adducts and triggering cell apoptosis, subpopulations of tumor cells can survive this stress (Pogge von Strandmann, Reinartz et al., 2017).

Tumor sequencing has revealed the complexity and heterogeneity of HGSOC (Cancer Genome Atlas Research, 2011). DNA breakage and regions of chromosomal gain or loss are common (Patch, Christie et al., 2015). Gains at 8q24 occur in the majority of HGSOC tumors and encompass the *MYC* oncogene at 8q24.21 (Gorringe, George et al., 2010). *MYC* facilitates pluripotent stem cell generation and contributes to HGSOC chemoresistance (Fagnocchi & Zippo, 2017, Kumari, Folk et al., 2017, Li, Bonazzoli et al., 2019). Although MYC expression is generally high in HGSOC, the clinical significance of this change remains unclear.

A major regulator of Myc protein expression is the Wnt/β-catenin signaling pathway, essential for embryonic development and activated in a variety of tumors (Shang, Hua et al., 2017). The Wnt and Myc signaling pathways are among the ten most prevalent oncogenic signaling pathways in cancer (Sanchez-Vega, Mina et al., 2018). Wnt signaling is tightly regulated by the stability, subcellular localization, and transcriptional activity of β-catenin supporting cancer stem cell (CSC) survival and chemoresistance (Condello, Morgan et al., 2015, Nagaraj, Joseph et al., 2015). Platinum treatment can, paradoxically, also enhance ovarian cancer ‘stemness’ (Wiechert, Saygin et al., 2016). Increased aldehyde dehydrogenase (ALDH) activity, arising from elevated expression of a family of cellular detoxifying enzymes, is a hallmark of ovarian CSCs (Raha, Wilson et al., 2014, Silva, Bai et al., 2011). The growth of cells as tumorspheres *in vitro* increases chemotherapy resistance, ALDH expression, de-differentiation and stemness (Malta, Sokolov et al., 2018, Shah & Landen, 2014). Notably, HGSOC spread involves peritoneal tumor dissemination and tumorsphere growth within ascites (Pogge von Strandmann et al., 2017).

The *PTK*2 gene at 8q24.3, encoding focal adhesion kinase (FAK), is frequently amplified in breast, uterine, cervical, and ovarian tumors (Kaveh, Baumbusch et al., 2016). FAK is a cytoplasmic tyrosine kinase canonically activated by integrin receptor signaling controlling cell movement (Mitra, Hanson et al., 2005). FAK auto-phosphorylation at tyrosine 397 (pY397) is a reporter of FAK activity (Kleinschmidt & Schlaepfer, 2017). HGSOC tumors with *PTK2* gains exhibit elevated FAK expression and FAK Y397 phosphorylation (Cancer Genome Atlas Research, 2011, Zhang, Liu et al., 2016). Metastatic HGSOC tumor micro-environments are enriched with matrix proteins that are activators of FAK (Pearce, Delaine-Smith et al., 2018). While FAK knockdown and FAK inhibitor studies support an important role in promoting invasive tumor growth (Tancioni, Uryu et al., 2014, Ward, Tancioni et al., 2013), the targets downstream of FAK are varied and may be tumor or stroma context-dependent (Haemmerle, Bottsford-Miller et al., 2016, Sulzmaier, Jean et al., 2014). Moreover, phenotypes associated with FAK knockout may be distinct from FAK inhibition, since kinase-inactive FAK retains important scaffolding functions (Lim, Chen et al., 2008).

Several ATP-competitive FAK inhibitors have been developed and have displayed acceptable Phase I safety profiles in patients with advanced solid tumors (Hirt, Waizenegger et al., 2018, Jones, Siu et al., 2015, Soria, Gan et al., 2016). Phase II trials with FAK inhibitors have yielded some complete responses and many patients exhibiting stable disease. Current combinatorial approaches with FAK inhibitors are being tested in patients with pancreatic, mesothelioma, and non-small cell lung carcinoma. In ovarian and prostate carcinoma mouse tumor models, FAK inhibition (VS-6063, defactinib) enhanced taxane-mediated tumor apoptosis (Kang, Hu et al., 2013, Lin, Lee et al., 2018). Additionally, inhibitors of FAK and Myc exhibit combinatorial activity in promoting HGSOC apoptosis *in vitro* (Xu, Lefringhouse et al., 2017). It remains uncertain whether gains in 8q24 encompassing *PTK2* are associated with specific HGSOC cell phenotypes or responses to therapy, as determinants of FAK pathway dependence in tumors remain unknown.

Herein, we molecularly characterize a new murine model of ovarian cancer that displays spontaneous gains in the <u>*K*</u>-*Ras*, <u>*M*</u>*yc*, and <u>F</u>AK genes among other striking similarities to human ovarian carcinoma cell phenotypes. By using a combination of genetic FAK knockout and rescue, pharmacological inhibition, sequencing and bioinformatics, we identify a non-canonical FAK activity-dependent linkage to β-catenin and Myc leading to differential mRNA target expression supporting CSC survival and platinum resistance. Our studies linking intrinsic FAK activity to platinum resistance support the combinatorial testing of FAK inhibitors for recurrent ovarian cancer.

## Results

### A new *in vivo* evolved murine epithelial ovarian cancer model

HGSOC is characterized by p53 mutation or loss and common genomic copy number alterations, yet no preclinical models exist to study connections between genomic gains or losses with cell phenotypes. Murine ID8 cells are spontaneously-immortalized clonal ovarian epithelial cells that can form tumors in normal C57Bl6 mice (Roby, Taylor et al., 2000). ID8 cells do not contain common oncogenic mutations, express wild type p53, and p53 inactivation enhances ID8 tumor growth and increases sensitivity to platinum chemotherapy (Walton, Blagih et al., 2016, Walton, Farquharson et al., 2017). Passage of ID8 cells through C57Bl6 mice can enhance ID8 tumorigenic potential via uncharacterized mechanisms (Clark, Gupta et al., 2016, Mo, Bachelder et al., 2015, Ward et al., 2013).

We previously isolated aggressive ID8-IP cells, lethal in mice within 40 versus 90 days compared to parental ID8 cells (Ward et al., 2013), via early recovery of ascites-associated cells and anchorage-independent expansion *ex vivo* (Fig. 1A). Total exome sequencing (90% of exons sequenced at 100X) of ID8 and ID8-IP cells revealed 19619 shared, 29373 ID8 unique, and 11800 ID8-IP unique gene variants (Supplemental Fig. S1 and Supplemental Tables S1-S3). However, less than 1% of exon variants identified were detected as mRNAs by RNA sequencing (~60 million clean reads/replicate). No murine-equivalent mutations were found in COSMIC, the Catalogue of Somatic Mutations in Cancer. In addition to ID8 non-synonymous mutations previously identified (Walton et al., 2016), we detected two additional potential damaging mutations in *HJURP* (Supplemental Fig. S1). In ID8-IP cells, after filtering for transcribed variants and subtracting for mutations found in the Single Nucleotide Polymorphism database (dbSNP138), predicted damaging mutations were identified in *XXYLT1* and *ATXN10* (Supplemental Fig. S1C). Overall, the mutational burden in ID8-IP cells is low.

**Figure 1.**
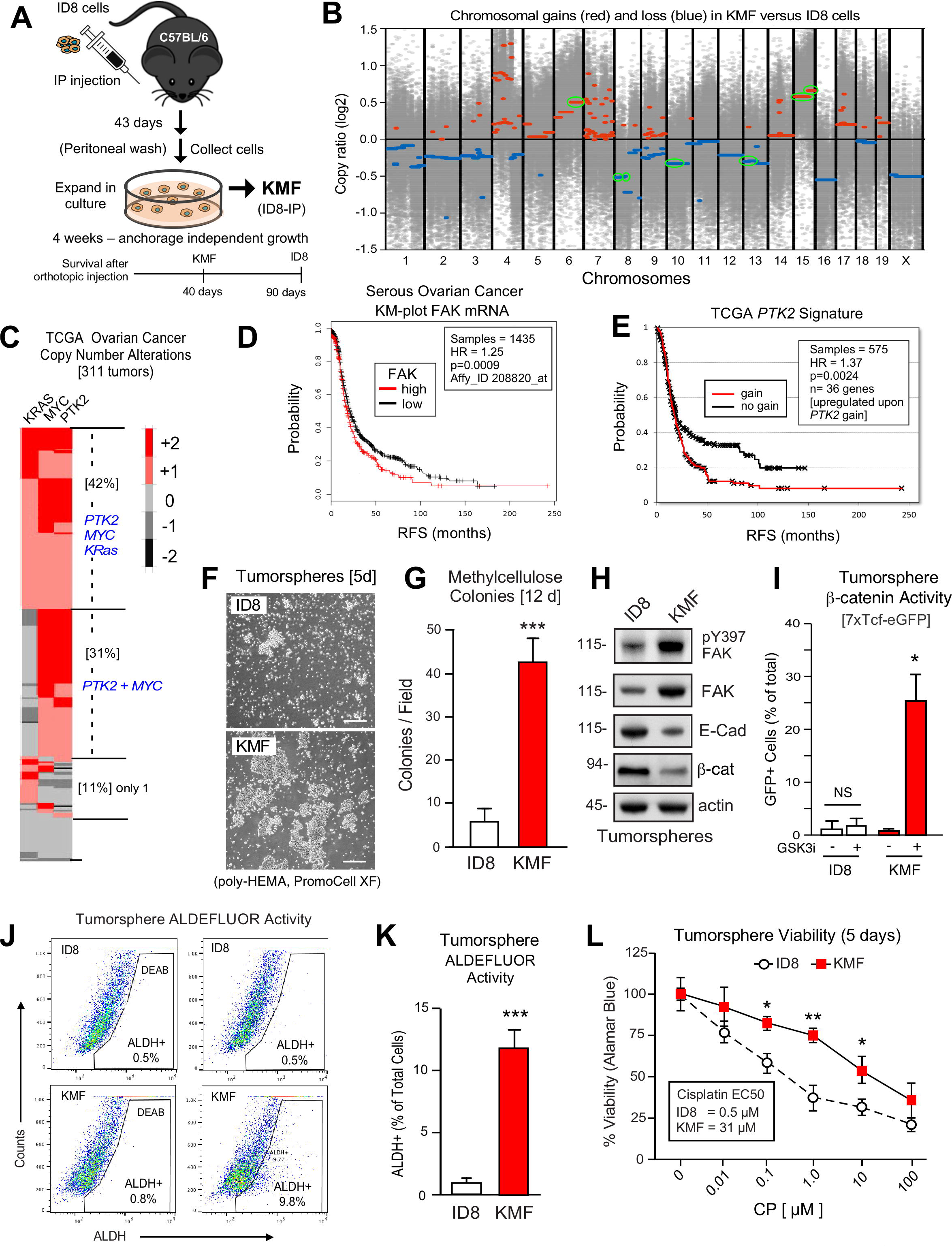
KMF (gains in genes for K-Ras, Myc, and FAK) murine model of ovarian cancer with shared characteristics to HGSOC. **(A)** Schematic of KMF cell isolation by in vivo selection for aggressive ID8 growth in C57Bl6 mice and expansion as tumorspheres. **(B)** Whole-genome copy number ratio (log2) determined from ID8 and KMF exome sequencing. Gains (red) and losses (blue) are denoted across chromosomes. Circled regions (green) highlight shared genomic copy alterations between KMF and HGSOC (Table 1). **(C)** Heat map showing genomic copy number alterations encompassing *KRAS*, *MYC*, and *PTK2* (FAK) genes in HGSOC patients (TCGA, 311 tumors). **(D)** Kaplan-Meier (KM) analysis of FAK mRNA levels in 1435 patient samples. Plot shows probability of relapse-free survival (RFS) in months with tumors high (red) or low (black) for FAK mRNA (HR = 1.25, P = 0.0009). **(E)** KM plot of *PTK2* signature (red, Supplemental Table S5) versus no gain (black) showing RFS (HR = 1.37, P = 0.0024). **(F)** Representative images of ID8 and KMF tumorsphere formation (5 days) in serum-free PromoCell XF media. Scale is 1 mm. **(G)** Quantitation of ID8 and KMF colony formation in methylcellulose (21 days). Values are means (± SEM, *** P <0.001, unpaired T-test) from 3 independent experiments. **(H)** ID8 and KMF 3D protein lysates immunoblotted for pY397 FAK, FAK, E-cadherin, β-catenin, and actin. **(I)** β-catenin transcriptional reporter activity (7X TCF-eGFP) in ID8 and KMF cells grown as tumorspheres +/− GSK3β inhibitor. Values are percent GFP+ cells by flow cytometry (NS, not significant, * P <0.05, unpaired T-test, 2 experiments). **(J)** Representative flow cytometry profiles of ID8 and KMF tumorspheres analyzed for ALDEFLUOR activity. N,N-diethylaminobenzaldehyde (DEAB) treatment was used to set gates for ALDH-bright (ALDH+) cells. **(K)** Quantitation of ID8 and KMF tumorsphere ALDEFLUOR activity. Values are means expressed as fold-change to ID8 (± SD, *** P <0.001, unpaired T-test, 3 independent experiments). **(L)** Tumorsphere cytotoxicity (Alamar Blue) with increasing CP (5 days) and expressed as percent viability to DMSO control. Means (n = 2) from 4 independent experiments (± SEM, * P <0.05, ** P < 0.01 by two-way ANOVA with a Bonferroni’s multiple comparisons test). EC50 values independently determined using Graphpad Prism.

To determine if gene copy number alterations underlie ID8-IP phenotypes, exome sequencing read values and bioinformatic analyses were used to map sites of DNA gains or loss across chromosomes using ID8 as a reference (Fig. 1B). Gains in murine chromosome cytoband regions 6qD1-G3, 15qD3-F3, and 15qA1-D3 were present in ID8-IP cells (Figure 1B, green circles). These correspond to human cytobands 12p12.1, 8q24.2, and 8q24.3, with the latter two representing one of the most amplified regions in HGSOC (Cancer Genome Atlas Research, 2011, Li et al., 2019). The gain in cytoband 15qA1-D3 is in addition to chromosome 15 polyploidy detected by ID8 karyotype analyses (Roby et al., 2000). Notably, common gene gains in ID8-IP and HGSOC include *KRAS*, *MYC*, and *PTK2* (encoding FAK) that support proliferation, stem cells, and adhesion signaling, respectively (Table I). Herein, these ID8-IP cells will be termed KMF to denote gains in <u>**K**</u>-Ras, <u>**M**</u>yc, and <u>**F**</u>AK genes. Murine KMF cells contain several gains or losses in genes common to the top 20 set of genes altered in HGSOC (Table I and Supplemental Table S4).

**Table 1.** Shared genetic alterations between murine KMF cells and human HGSOC

| Murine Cytoband | Human Cytoband | Gain/Loss | Genes in Common murine-human loci | Pathway/Role |
| --- | --- | --- | --- | --- |
| 6qD1-G3 | 12p12.1 | Gain | *KRAS* | Proliferation |
| 15qA1-D3 | 8q24.21, 8q24.3 | Gain | *MYC, PTK2* | Stem Cell, Adhesion |
| 15qD3-F3 | 8q24.3 | Gain | *RECQL4* | DNA Repair |
| 8qA1.1-1.3 | 8p23.3 | Loss | *TUSC3* | Tumor Suppressor |
| 8qB1.1-1.2 | 4q34.3 | Loss | *IRF2* | Interferon Response |
| 10qA1-D1 | 19p13.3 | Loss | *TJP3* | Tight Junction |
| 13qB3-D2.3 | 5q11.2, 5q13.1 | Loss | *MAP3K1, FOXD1, PIK3R1* | MAPK, Cell Cycle<br>P85 (PI3-kinase adaptor) |

Shared copy number alterations with top 20 most significant amplifications and deletions in HGSOC (Cancer Genome Atlas Research, 2011)

### MYC and PTK2 associations in HGSOC

In HGSOC, *KRAS*, *MYC*, and *PTK2* gains co-occur in 42% of tumors and *PTK2* plus *MYC* co-occur in an additional 32% of HGSOC patients (Fig. 1C). More than 70% of HGSOC tumors contain combined gains at *PTK2* and *MYC* loci. Although MYC has oncogenic properties (Kumari et al., 2017), MYC copy number gains are not strongly correlated with increased MYC mRNA or protein levels in HGSOC patient tumors (Supplemental Figs. S2). By contrast, *PTK2* copy number gains are linearly proportional to increased FAK mRNA (R^2^ = 0.66) and FAK protein (R^2^ =0.61) levels in HGSOC (Supplemental Figs. S2). Elevated FAK mRNA levels are associated with decreased HGSOC patient relapse-free survival (n = 1435, P = 0.0009, and hazard ratio = 1.25) (Fig. 1D). Bioinformatic analyses identified a set of 36 genes on different chromosomes in HGSOC that exhibit a significant and at least a two-fold change in tumors with elevated *PTK2* (Supplemental Table S5). This 36 gene set was associated with a significant shorter time to relapse (n = 575, P = 0.0024, and hazard ratio = 1.37) (Fig. 1E). Together, these results support the importance of *PTK2* gains, and indirectly FAK protein, in promoting HGSOC tumor progression.

### KMF cells exhibit enhanced CSC phenotypes and cisplatin resistance

KMF cells also contain regions of chromosome loss that correspond to regions frequently eliminated in HGSOC encompassing *GNA11, MAP2K2, TJP3, MAP3K1, CCNB1, PIK3R1*, and *FOXD1* genes (Table 1). Additionally, a mutation in *TRP53* exon 11 encoding the 3’ untranslated region and two p53 mRNA splice variants were detected within KMF cells (Supplemental Fig. S3). Immunoblotting did not detect p53 protein expression in KMF cells compared to ID8 and murine Lewis lung carcinoma cells containing wildtype p53. Several p53 target mRNAs were decreased in KMF compared to ID8 cells under 3D conditions by quantitative real-time PCR (Supplemental Fig. S3). Surprisingly, KMF cells exhibit constitutively-elevated p21CIP1 protein, a known target of p53 transcriptional activity (Supplemental Fig. S3). However, deregulated p21CIP1 levels can be indicative of a p53-deficient environment (Georgakilas, Martin et al., 2017). RNA sequencing did not detect expression of Cdkn2a (p16/INK4A) or Cdkn2b (p15/INK4B), which impact p53 protein stability and cell transformation (Al-Khalaf & Aboussekhra, 2018). We conclude that KMF cells are *de facto* deficient in p53 regulation.

KMF cells possess a mesenchymal-like morphology compared to the epithelial-like growth of ID8 cells in 2D culture (Supplemental Fig. S4). KMF cells form more 3D tumorspheres (Fig. 1F) and colonies in methylcellulose than ID8 cells (Fig. 1G). RNA sequencing revealed 10800 shared, 744 ID8 enriched, and 402 KMF enriched transcripts with FPKM (Fragments Per Kilobase of transcript per Million mapped read) values > 1 (Supplemental Fig. S4 and Supplemental Table S6). KMF tumorspheres exhibit target enrichment in *Cell Cycle*, *Mitotic Spindle Checkpoint*, and *DNA Repair* pathways (Supplemental Figs. S4). At the protein level, KMF tumorspheres expressed elevated FAK, increased FAK pY397, but decreased E-cadherin and β-catenin levels relative to ID8 cells (Fig. 1H). Although KMF cells expressed lower β-catenin protein levels, β-catenin transcriptional activity was greatly elevated in KMF compared to ID8 cells grown as tumorspheres (Fig. 1I). As FAK and β-catenin are co-regulators of CSC phenotypes in breast carcinoma cells (Kolev, Tam et al., 2017), ID8 and KMF cells were analyzed for changes in ALDH activity and intrinsic CP resistance. KMF tumorspheres exhibited increased ALDH activity (Figs. 1J and K) and were more resistant to cisplatin (CP)-induced toxicity than ID8 cells (Fig. 1L). The enhanced CSC phenotypes and intrinsic CP chemoresistance of KMF cells remained after extended passage. Taken together, this new KMF model exhibits noteworthy similarities to HGSOC.

### FAK pY397 remains elevated in ovarian carcinoma tumors surviving neoadjuvant chemotherapy

A subset of HGSOC patients are treated with neoadjuvant carboplatin and paclitaxel chemotherapy to reduce tumor burden prior to surgery (Matulonis et al., 2016). However, some tumor cells, such as CSCs, escape CP-mediated apoptosis and survive chemotherapy (Wiechert et al., 2016). FAK protein and FAK tyrosine phosphorylation (pY397 FAK) levels are elevated in primary HGSOC tumors compared to normal tissue (Zhang et al., 2016), but it is not known whether chemotherapy alters FAK activation. To evaluate this, paired primary biopsies and tumors obtained at the time of cytoreductive surgery following neoadjuvant carboplatin and paclitaxel chemotherapy were analyzed by immunohistochemical (IHC) staining and quantitative image analyses (Supplemental Table S3 and Supplemental Fig. S5). A high degree of Pax8 and pY397 FAK co-localized staining was detected (Fig. 2A) with FAK pY397 exhibiting both cytoplasmic and nuclear localization. Many of these tumor cells were positive for the Ki-67 proliferation marker. Validation was obtained using a pY397 FAK peptide block (Supplemental Fig. S5) and FAK pY397 staining was objectively higher in ovarian tumor compared to stromal cells (Fig. 2B). Among tissue samples obtained after neoadjuvant chemotherapy, Pax8 and pY397 FAK staining remained highly co-localized within residual tumors (Fig. 2C and Supplemental Fig. S5). Surprisingly, by comparing samples from the same patients, pY397 FAK staining after neoadjuvant chemotherapy in residual tumor cells was significantly increased (Fig. 2D).

**Figure 2.**
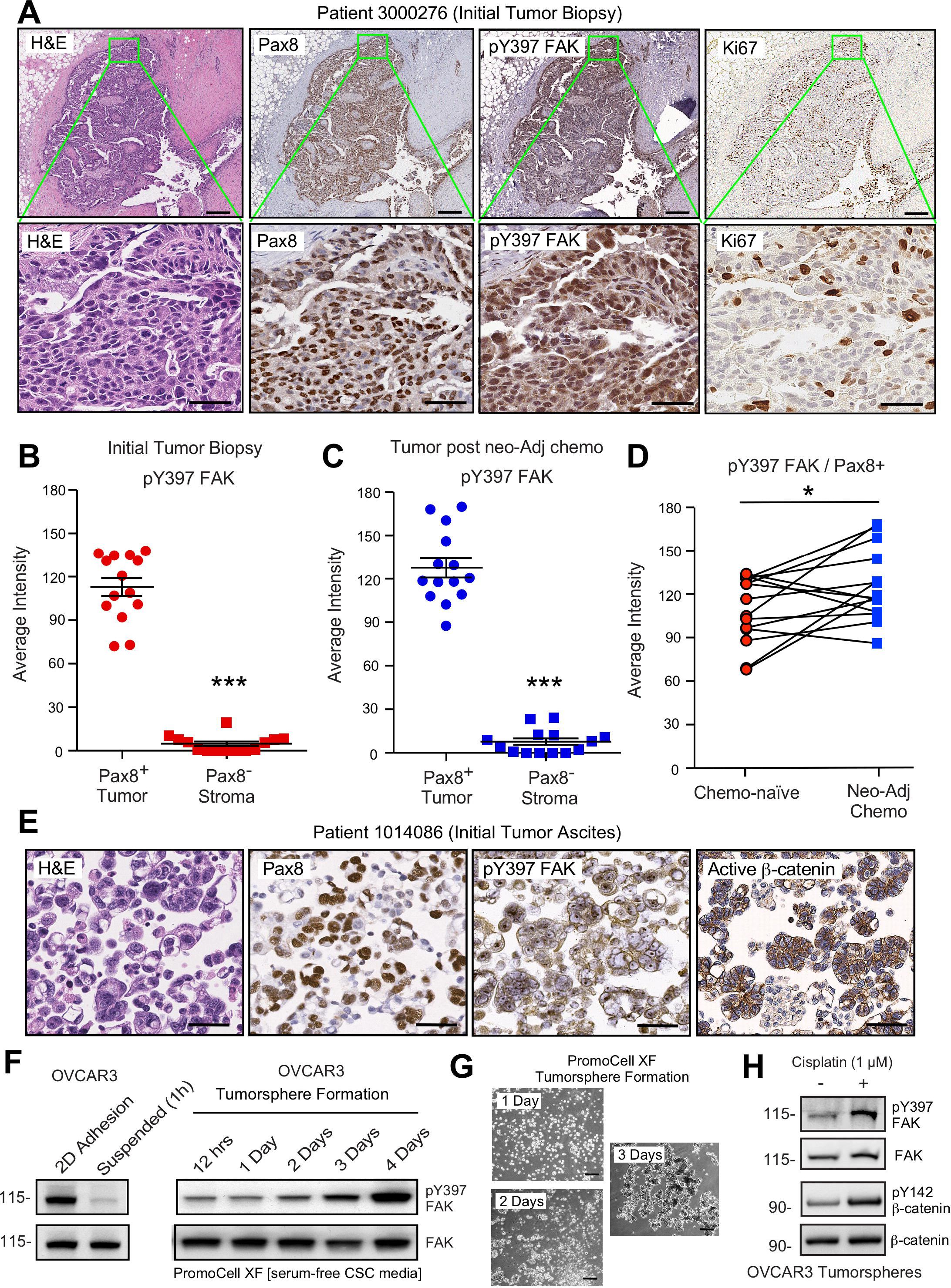
Elevated FAK Y397 phosphorylation (pY397) in HGSOC patient tumors surviving neoadjuvant chemotherapy. **(A)** IHC staining of paraffin-embedded serial sections (patient 3000276, Supplemental Table S7) with H&E, Pax8, pY397 FAK, and Ki67. Scale is 200 µm. Inset (green box) region is shown at 40X (below). Scale is 60 µm. **(B** and **C)** FAK pY397 staining intensity of paired patient ovarian tumor samples from initial biopsies (panel B) and after surgical removal following neoadjuvant chemotherapy (panel C) within Pax8-positive (tumor) and Pax8-negative (stroma) regions. Dot plots are quantified staining from 14 paired patient samples (Aperio software) and bars show mean ± SEM (analyzed 11 regions per sample, *** P <0.001, unpaired T-test). **(D)** Increased FAK pY397 staining within Pax8-positive regions post-chemotherapy (* P <0.05, paired T-test). **(E)** IHC serial section staining (H&E, Pax8, pY397 FAK, and active β-catenin) of peritoneal ascites cells (tumorspheres) from initial patient biopsy. **(F)** OVCAR3 lysates from 2D adherent, suspended (1 h), and cells in anchorage-independent serum-free (PromoCell XF) conditions facilitating tumorsphere formation were analyzed by total FAK and pY397 FAK immunoblotting. **(G)** Representative images of OVCAR3 tumorsphere formation. Scale is 2 mm. **(H)** OVCAR3 cells as tumorspheres (Day 3) treated with DMSO or CP (1 µM) for 1 h and protein lysates blotted for pY397 FAK, total FAK, pY142 β-catenin, and total β-catenin.

Increased FAK pY397 is generally considered a marker for elevated cell adhesion or tissue stiffness (Sulzmaier et al., 2014). Unexpectedly, elevated pY397 FAK staining was also observed within ascites Pax-8 positive tumorspheres that displayed active β-catenin staining (Fig. 2E). This was unanticipated, since FAK Y397 phosphorylation is rapidly lost when human platinum resistant OVCAR3 cells are removed from adherent 2D culture and held in suspension (Fig. 2F). However, an extended time course analyses of OVCAR3 cells placed in anchorage-independent defined media conditions (PromoCell XF) revealed that FAK Y397 phosphorylation was restored as OVCAR3 cells clustered to form tumorspheres within 2-3 days (Figs. 2F and G). Surprisingly, CP (1 µM) treatment of OVCAR3 tumorspheres (EC50 > 10 µM) also resulted in increased FAK Y397 and β-catenin Y142 phosphorylation (Fig. 2H). As β-catenin Y142 is a direct FAK substrate promoting β-catenin activation in endothelial cells (Chen, Nam et al., 2012), our findings support the notion that non-canonical FAK activation occurs during tumorsphere formation and in response to CP stimulation.

### Combinatorial effects of CP and FAK inhibition

As CP treatment increased FAK Y397 phosphorylation, and since FAK activation was associated with more aggressive phenotypes in patient tumors and in KMF cells, we next investigated the effect of low dose CP treatment (1 µM) in the presence or absence of FAK inhibition (VS-4718, 1 µM) over 5 days on tumorsphere formation, ALDH activity, and viability of OVCAR3 and KMF cells (Figs. 3A-C). Cisplatin EC50 values for growth inhibition were 13 µM and 31 µM for OVCAR3 and KMF tumorspheres, respectively. CP treatment resulted in increased tumorsphere formation and ALDEFLUOR activity of OVCAR3 and KMF cells, consistent with this being an activation-type stress. In contrast, FAK inhibitor (FAKi or VS-4718, 1 µM) treatment significantly reduced tumorsphere formation and ALDEFLUOR activity compared to control OVCAR3 and KMF cells (Figs. 3A and B).

**Figure 3.**
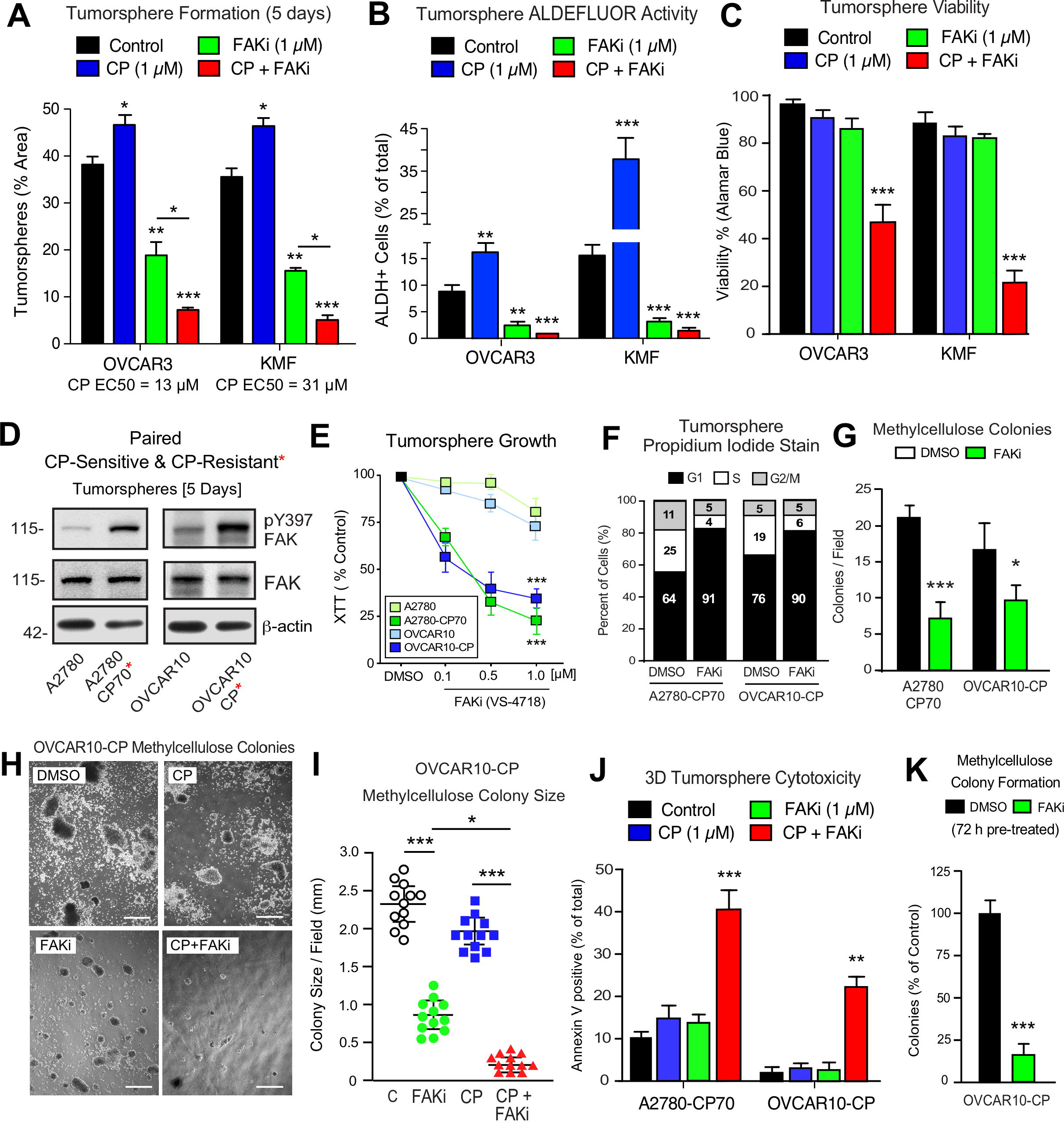
Prevention of CSC phenotypes *in vitro* by pharmacological FAK inhibition. **(A-C)** Quantification of OVCAR3 and KMF tumorsphere formation (panel A), ALDEFLUOR activity (panel B), and tumorsphere viability (panel C) in the presence of DMSO (control), CP (1 µM), FAKi (VS-4718, 1 µM) or CP plus FAKi. Values are means (± SEM, * P <0.05, ** P <0.01, *** P < 0.001 unpaired T-test) of 3 independent experiments. Panel C, values are means (± SEM, *** P <0.001, one-way ANOVA) from 3 independent experiments. **(D)** Human A2780, A2780-CP70, OVCAR10, and OVCAR10-CP tumorsphere lysates immunoblotted for FAK pY397, FAK, and actin. **(E)** Growth of A2780, OVCAR10, A2780-CP70, or OVCAR10-CP cells as tumorspheres in the presence of FAKi (VS-4718, 0.1 to µM) for 4 days. Values are means (± SEM, *** P <0.001, one-way ANOVA) from 2 independent experiments. **(F)** A2780-CP70 or OVCAR10-CP cells grown as tumorspheres (3 days) were treated with DMSO or FAKi (VS-4718, 1 µM) for 24 h, stained with propidium iodide, and analyzed by flow cytometry. Percentage of cells in G0/G1, S, or G2-M phase of the cell cycle determined using FlowJo. **(G)** Quantitation of A2780-CP70 and OVCAR10-CP colony formation in methylcellulose (21 days) with DMSO (control) or FAKi (VS-4718, 1 µM). Values are means (± SEM, *P<0.05, *** P <0.001, unpaired T-test) from 2 independent experiments. **(H and I)** Representative OVCAR10-CP methylcellulose colony formation (21 days) (panel H) and colony size determination (panel I) in the presence of DMSO (control), CP (1 µM), FAKi (1 µM), or CP+FAKi. Scale is 2.5 mm. Values are means (± SEM, * P<0.05, *** P<0.001, one-way ANOVA) from 2 independent experiments. **(J)** A2780-CP70 and OVCAR10-CP tumorsphere cytotoxicity (annexin V) in the presence of DMSO (control), CP (1 µM), FAKi (1 µM), or CP+FAKi. Values are means (± SEM, ** P<0.01, one-way ANOVA) from 3 independent experiments. **(K)** Secondary OVCAR10-CP colony formation after pre-treatment with FAKi (1 µM, 72 h in 2D conditions). Values are means (± SEM, *** P <0.001) from 2 independent experiments.

FAKi was not directly cytotoxic, since only CP combined with FAKi reduced tumorsphere viability (Fig. 3C). Single agent CP or FAKi treatment did not alter KMF (Supplemental Fig. S6) or OVCAR3 (Supplemental Fig. S7) growth or viability in 2D culture. FAKi treatment inhibited FAK Y397 phosphorylation in 2D culture, but this did not impact 2D cell growth. Under 3D conditions, FAKi similarly reduced FAK Y397 phosphorylation, and was associated with an elevated percentage of KMF and OVCAR3 cells in G1 phase of the cell cycle, but alone did not trigger apoptosis (Supplemental Figs. S6 and S7). The finding that FAKi decreased 3D, but not 2D cell proliferation, and that FAKi exhibits combinatorial activity with low-dose CP to promote cell apoptosis is consistent with a non-canonical role of context-dependent FAK signaling supporting tumorsphere growth and survival.

### Constitutive FAK activity is connected to acquired CP-resistant phenotypes

Phosphoinositide 3-kinase (PI3K)-elicited Akt activation is one of several survival signaling pathways downstream of FAK (Sulzmaier et al., 2014). More than half of HGSOC harbor genetic lesions that can elevate PI3K activity (Hanrahan, Schultz et al., 2012). A2780 human ovarian carcinoma tumor cells contain activating mutations in *PI3KCA* and inactivation of *PTEN* - alterations that can promote Akt activation (Domcke, Sinha et al., 2013). OVCAR10 cells similarly exhibit elevated Akt phosphorylation. Both A2780 and OVCAR10 cells are resistant to FAKi (1 μM) effects on 3D cell proliferation (Tancioni et al., 2014). To determine if *in vitro* acquisition of increased CP resistance alters responses to FAKi, intermittent CP exposure (10 μM) and cell recovery was used to generate OVCAR10-CP and maintain A2780-CP70 cells (Godwin, Meister et al., 1992). Both A2780-CP70 (EC50 = 60 μM) and OVCAR10-CP (EC50 = 9 μM) cells maintain elevated CP resistance and immunoblotting revealed constitutively-elevated FAK pY397 within tumorspheres of CP-resistant cells compared to parental cells (Fig. 3D). Unexpectedly, CP-resistant A2780-CP70 and OVCAR10-CP cells exhibited a newly-acquired dose-dependent sensitivity to FAKi growth inhibition as tumorspheres (Fig. 3E), but not when the same cells were grown in 2D conditions (Supplemental Fig. S8).

Pharmacological FAK inhibition of A2780-CP70 and OVCAR10-CP tumorsphere growth was accompanied by an increased number of cells in G1 phase of the cell cycle (Fig. 3F), decreased cyclin D1 expression, but not increased apoptosis (Supplemental Fig. S8). FAKi prevented A2780-CP70 and OVCAR10-CP methylcellulose colony formation (Fig. 3G), but FAKi had no effect on parental A2780 or OVCAR10 colony formation (Supplemental Fig. S8). An alternate FAKi (VS-6063, defactinib) also prevented A2780-CP70 colony formation in a dose-dependent manner (Supplemental Fig. S9). Both A2780-CP70 and OVCAR10-CP tumorspheres exhibited increased ALDH activity compared to parental cells and this was dependent on FAK activity (Supplemental Fig. S10).

OVCAR10-CP colony formation was evaluated in the presence of DMSO (control), FAKi, CP, or FAKi and CP to evaluate whether acquired sensitivity to FAKi could impact responses to CP treatment (Fig. 3H). Single agent FAKi reduced colony size (Fig. 3I), consistent with an inhibitory effect on tumorsphere proliferation (Figs. 3E and F). The combination of FAKi with CP prevented colony formation (Figs. 3H and I) and was associated with increased OVCAR10-CP apoptosis (Fig. 3J). Increased A2780-CP70 apoptosis also occurred only in the presence of FAKi and CP (Fig. 3J). As OVCAR10-CP cells pre-treated with FAKi had a reduced ability to form methylcellulose colonies (Fig. 3K), our results point to the importance of FAK activity in promoting elevated ALDH levels and CSC phenotypes in *PI3KCA-* and *PTEN*-mutated A2780-CP70 and OVCAR10-CP cells as part of adaptive changes associated with acquired CP resistance.

### FAK inhibition prevents KMF CSC survival *in vivo*

ALDH activity and secondary tumor initiation frequency (TIF) are reporters used to identify CSCs (Raha et al., 2014). Previous studies with triple-negative breast carcinoma models showed that FAKi (VS-4718) triggered CSC apoptosis (Kolev et al., 2017). As FAKi treatment prevents KMF tumorsphere associated ALDH activity independent of apoptosis (Fig. 3), we next tested KMF secondary TIF capability after treating KMF tumor-bearing mice with VS-4718 (50 mg/kg, BID) for 21 days (Figs. 4A-D). Tumor formation and the presence of mCherry-labeled KMF cells in peritoneal ascites were reduced by FAKi (Supplemental Fig. S11). FAK Y397 phosphorylation was similarly suppressed in ovarian tumors in response to oral FAKi administration (Fig. 4B). ALDH activity was detected in a high percentage of peritoneal wash-collected cells from vehicle-treated mice and ALDH-positive cells were significantly reduced *in vivo* by FAKi treatment (Fig. 4C and Supplemental Fig. S11). KMF cells re-isolated from vehicle treated mice possessed high secondary TIF, forming tumors in 75% of mice injected with 300 cells (Fig. 4D). Importantly, viable KMF cells re-isolated from FAKi treated mice exhibited an approximately 85-fold reduction in TIF, with 12.5% of mice forming secondary tumors upon injection of 2000 cells (Fig. 4D). These results support the notion that FAK inhibition compromises CSCs *in vivo* and critically, that FAKi reversed *de novo* gains in KMF CSC phenotypes.

**Figure 4.**
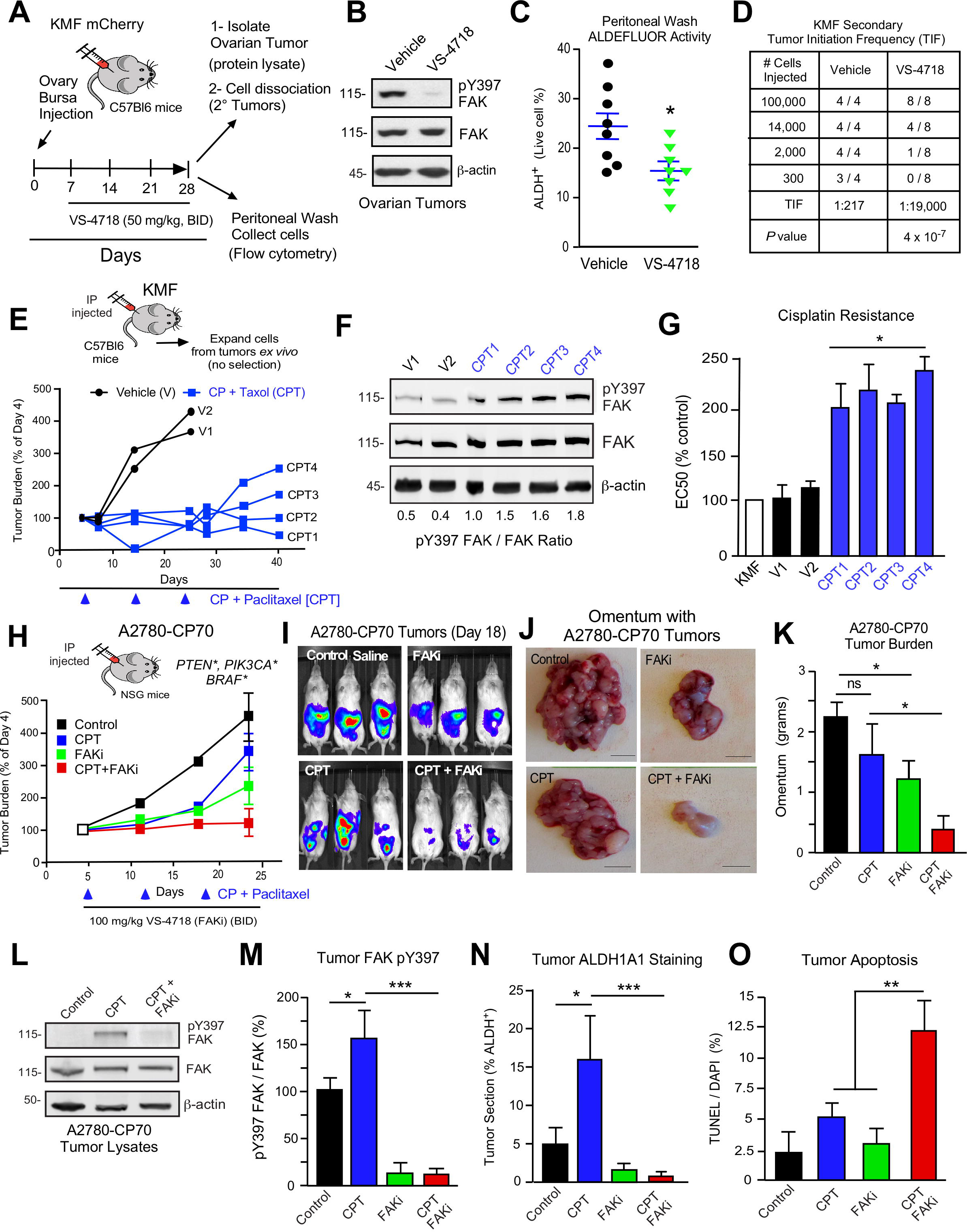
FAK activity supports *in vivo* CSC survival and pharmacological FAK inhibition sensitizes CP-resistant tumors chemotherapy-induced apoptosis. **(A)** Experimental schematic for evaluation of mCherry-KMF orthotopic tumors in vehicle- or VS-4718-treated mice. **(B)** KMF tumor lysates immunoblotted for pY397 FAK, FAK and actin. **(C)** Percent of KMF ALDH+ cells collected in peritoneal wash from vehicle or VS-4718-treated mice (n=8 mice each, ± SEM, * P < 0.05, T-test). **(D)** CSC-associated secondary tumor initiation frequency (TIF) (7 weeks) using a dilution series of KMF cells isolated from vehicle or VS-4718-treated mice. **(E)** Experimental schematic and IVIS imaging of labeled KMF tumor burden after cisplatin (CP) plus paclitaxel treatment at Days 7, 14, and 21. Values are individual mice (percent of Day 4) and on Day 24 (V1 and V2) or Day 42 (CPT1-4). **(F)** Lysates of KMF cells isolated from vehicle or CPT-treated mice immunoblotted for FAK pY397, FAK, and actin. Mean LiCor quantitation of FAK pY397 to total FAK ratios (n=2 independent experiments) are shown. **(G)** Cisplatin EC50 values of ex vivo-expanded KMF cell (V1, V2, CPT1-CPT4) growth as determined by XTT assay. Values are means (± SEM, * P < 0.05, one-way ANOVA). **(H)** Experimental schematic and IVIS imaging of labeled A2780-CP70 cells IP injected into NSG mice (randomized at Day 5). Experimental groups: control saline (black) injection on Days 5, 12, and 19; VS-4718 by oral gavage (green, 100 mg/kg, BID); CPT chemotherapy injection (blue, 3 mg/kg cisplatin and 2 mg/kg paclitaxel) on Days 5, 12, and 19; and VS-4718 plus CPT combined administration (red). IVIS imaging was performed on Days 4, 11, 18, and 23. Tumor burden is expressed as percent of Day 4. **(I)** Representative IVIS images of A2780-CP70 tumor burden on Day 18. **(J)** Representative images of omentum with A2780-CP70 tumors at Day 24. Scale is 0.5 cm. **(K)** Omentum-associated A2780-CP70 tumor mass (n=6, ± SEM * P < 0.05, one-way ANOVA) from each treatment group. **(L)** A2780-CP70 tumor lysates immunoblotted for FAK pY397, FAK, and actin. **(M)** Ratio of pY397 FAK to total FAK levels in tumor lysates by immunoblotting. Values are means (± SEM * P < 0.05, *** P < 0.001, one-way ANOVA) of 3 tumors per experimental group. Control set to 100. **(N** and **O)** Percent ALDH-1A1 positive immunofluorescent A2780-CP70 tumor staining or apoptosis (TUNEL and Hoescht 33342 staining) in A2780-CP70 tumors. Values are means (± SEM, 2 independent tumors, 5 random fields per tumor at 20X, * P <0.05, ** P <0.01, *** P <0.001 one-way ANOVA).

### Increased FAK pY397 and CP resistance after chemotherapy

KMF tumorspheres exhibited increased FAK Y397 and elevated intrinsic CP resistance (Fig. 1). After carboplatin and paclitaxel neoadjuvant chemotherapy, HGSOC patient tumors similarly exhibited elevated FAK Y397 phosphorylation (Fig. 2). KMF tumor bearing mice were treated with CP and paclitaxel (CPT) chemotherapy to determine if elevated FAK Y397 phosphorylation after chemotherapy was associated with increased CP resistance. This treatment acted to limit, but did not prevent, KMF tumor growth (Fig. 4E). Re-isolation and expansion of peritoneal KMF cells from vehicle (V) or CPT treated mice followed by immunoblotting (Fig. 4F) revealed an increase increased pY397 FAK within KMF cells from tumor bearing mice that received CPT relative to vehicle treated control. Moreover, KMF cells from CPT-treated mice exhibited elevated CP resistance *in vitro* (Fig. 4G). Taken together, these results support a link between increased FAK pY397 and *in vivo* acquired CP resistance.

### FAK inhibition re-sensitizes CP resistant A2780-CP70 tumors to CPT chemotherapy

DTomato plus luciferase-labeled A2780 or A2780-CP70 cells were orthotopically injected into mice to assess the combinatorial potential of FAKi (VS-4718) and CPT chemotherapy on paired CP sensitive and CP resistant tumors. After tumors were established, CPT or FAKi, or both, were used as chemotherapy. A2780 tumor growth was insensitive to FAKi (Supplemental Fig. S12), consistent with limited FAKi effects on A2780 growth *in vitro*. In dramatic contrast, single-agent FAKi treatment reduced A2780-CP70 tumor growth approximately 40% compared to controls (Figs. 4H-K), suggesting that these CP resistant tumors acquire a FAK dependence. Flow cytometry analyses of peritoneal dTomato-labeled cells of tumor-bearing mice revealed that FAKi treatment selectively decreased ALDH activity in A2780-CP70, but not A2780, cells *in vivo* (Supplemental Fig. S13). This parallels the results obtained in the tumorsphere models (Fig. 3) and confirms that A2780-CP70 tumor growth, and signals regulating ALDH activity, are FAK dependent.

As expected, the growth of platinum-sensitive A2780 tumors is prevented by CPT chemotherapy (Supplemental Fig. S12), whereas final A2780-CP70 tumor burden after CPT treatment was only slightly less than untreated controls (Figs. 4H-K). Interestingly, CPT treatment increased FAK Y397 phosphorylation (Figs. 4L and M) and ALDH-1A1 positive staining (Fig. 4N) in non-necrotic regions of A2780-CP70 tumors (Supplemental Fig. S14). We hypothesized that gains in pY397 FAK and ALDH after CPT treatment may reflect FAK-dependent A2780-CP70 tumor cell survival signaling. Testing this hypothesis, we found that adding FAKi to CPT chemotherapy prevented A2780-CP70 tumor growth concurrent with suppression of FAK Y397 phosphorylation, loss of ALDH-1A1 expression, and tumor apoptosis *in vivo* (Figs. 4K-O). Together, these results show that FAK inhibition acts to re-sensitize CP-resistant A2780-CP70 tumors to CPT chemotherapy.

### PTK2 inactivation alters CSC phenotypes and CP resistance

As pharmacological FAK inhibition combined with CP resulted in CP-resistant tumor cell apoptosis *in vitro* and *in vivo*, we used CRISPR/Cas9 genetic targeting to introduce frameshift changes in human OVCAR3 *PTK2* exon 3 to create a FAK knockout (FAK-/-) model of HGSOC. Several FAK-/- clones were identified by immunoblotting and *PTK2* exon 3 inactivation was verified by DNA sequencing (Fig. 5A). Two clones (AB21 and AB28) were chosen for further characterization. OVCAR3-AB21 an -AB28 grew normally in 2D cell culture (Fig. 5B), but did not grow colonies in methylcellulose when seeded as single cells compared to parental OVCAR3 cells (Figs. 5B and C). However, when cultured in serum-free tumorsphere media over 5 days, parental OVCAR3 and OVCAR3 FAK-/- AB21 cells exhibited similar metabolic activity and viability (Fig. 5D).

**Figure 5.**
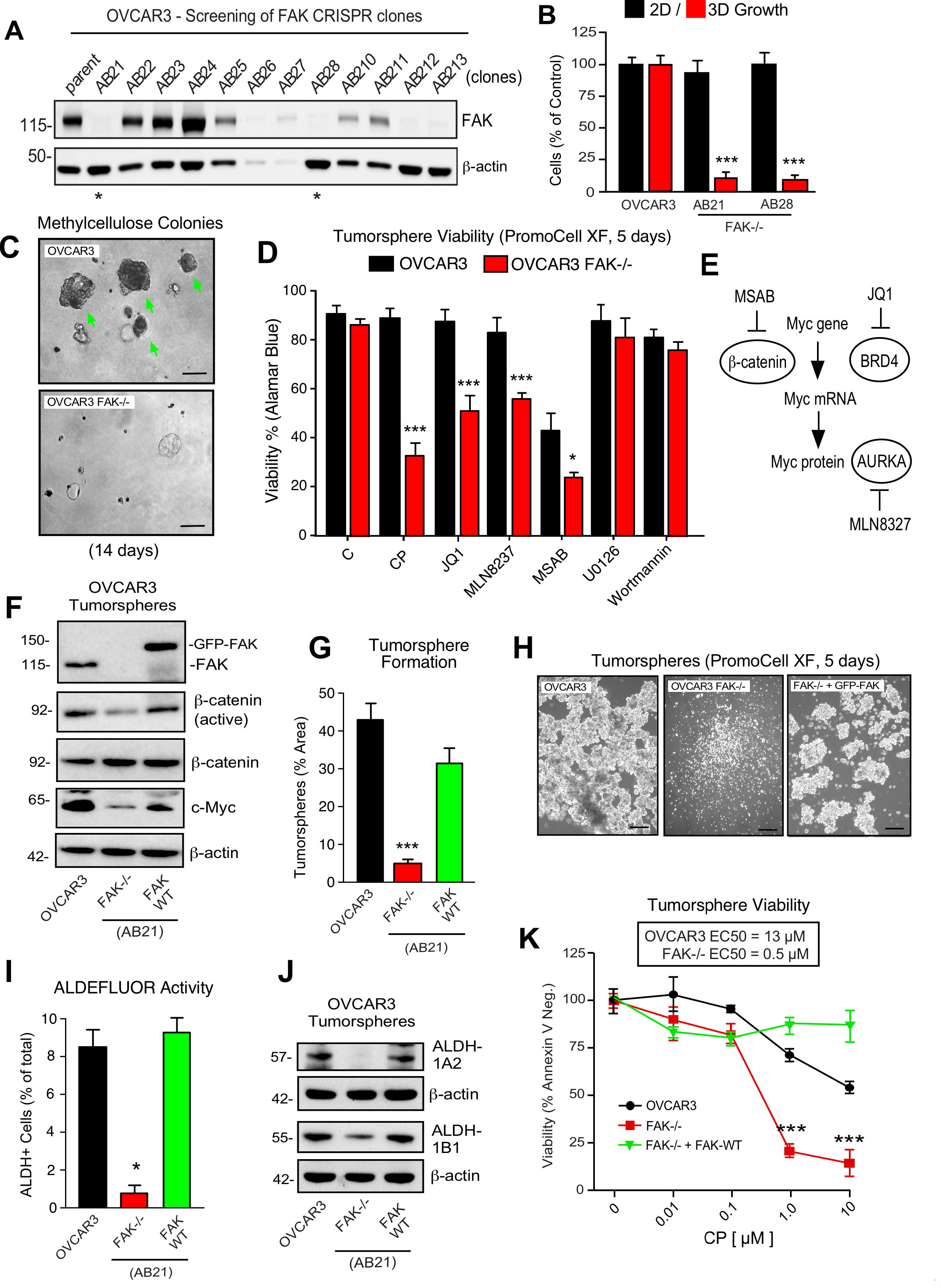
OVCAR3 CRISPR-mediated FAK knockout and re-expression. **(A)** Immunoblotting for FAK and actin in OVCAR3 lysates and identification of CRISPR-Cas9 FAK-/- clones (AB21 and AB28 are starred). **(B)** Growth of OVCAR3 parental, AB21, and AB28 cells in 2D culture (black bars) or enumeration of colonies in methylcellulose after 14 days (red bars). Values are means as percent of OVCAR3 control (± SEM, *** P <0.001, one-way ANOVA or NS, not significant) from 2 independent experiments. **(C)** Representative images of OVCAR3 (green arrows) or AB21 growth as colonies in merthylcellulose. Scale is 1 mm. **(D)** OVCAR3 or OVCAR3 FAK-/- (AB21) cell viability treated with DMSO (control), CP (1 μM), JQ1 (1 μM), MLN8237 (1 μM), MSAB (10 μM), U0126 (10 μM), or wortmannin (1 μM) after 72 h as measured by AlamarBlue. Values are means (± SEM, * P<0.05, *** P <0.001, one-way ANOVA with Fisher’s LSD multiple comparison test) for 3 independent experiments. **(E)** Schematic of signaling linkage of regulators of Myc gene expression (MSAB and JQ1) or Myc protein stability (MLN8327). **(F)** Immunoblotting for FAK, active β-catenin, β-catenin, Myc, and actin in lysates of OVCAR3, OVCAR3 FAK-/- (AB21), and OVCAR3 FAK-/- + GFP-FAK-WT cells in 3D conditions. **(G)** Tumorsphere formation (5 days). **(H)** Representative images of OVCAR3, OVCAR3 FAK-/- (AB21), and OVCAR3 FAK-/- + GFP-FAK-WT cells in PromoCell XF (5 days). Scale is 1 mm. **(I)** ALDEFLUOR activity. **(G** and **I)** Values are means (± SEM, n=2, *P <0.05, *** P <0.001, one-way ANOVA with a Tukey’s multiple comparisons test) from 3 (panel G) or 4 (panel I) independent experiments. **(J)** Immunoblotting for ALDH-1A2, ALDH-1B1, or actin in the indicated cell lysates. **(K)** Cytotoxicity (percent Annexin V negative) of OVCAR3 (black circles), OVCAR3 FAK-/- (red squares), and OVCAR3 FAK-/- + GFP-FAK-WT (green triangles) cells in PromoCell XF treated with increasing CP concentrations for 5 days. Values are means (± SEM, *** P <0.001, two-way ANOVA with a Bonferroni’s multiple comparisons test) from 3 independent experiments. EC50 values were determined independently and calculated using Prism (Graphpad).

Sustainability of OVCAR3 FAK-/- AB21 cells in 3D culture permitted a comparative evaluation of OVCAR3 cell sensitivities to different pharmacological inhibitors in a FAK-dependent manner. As measured by a fluorescent indicator of aerobic cell respiration, FAK-/- cell viability was inhibited in the presence of low dose CP, the bromodomain inhibitor JQ1, and the Aurora A kinase inhibitor MLN8237, but not the MEK1 inhibitor U0126 or the PI-3’kinase inhibitor wortmannin (Fig. 5D). JQ1 and FAKi can exhibit combinatorial inhibitory effects (Xu et al., 2017), while Aurora A can function to promote ovarian carcinoma adhesion and migration (Do, Xiao et al., 2014); pathways notably associated with FAK activation (Kleinschmidt and Schlaepfer, 2017). FAK-/- OVCAR3 cells exhibited increased cytotoxicity to methyl 3-(4-methylphenyl) sulfonyl amino-benzoate (MSAB), a small molecule promoting β-catenin degradation and inhibition of Wnt-dependent tumors (Hwang, Deng et al., 2016) (Supplemental Fig. S15). These results support the importance of β-catenin in OVCAR3 tumorsphere survival. Moreover, genetic FAK inactivation fosters sensitivity to pharmacological agents that target signaling pathways commonly impacting Myc gene expression or Myc protein stability (Fig. 5E) (Chen, Liu et al., 2018).

As AB21 FAK-/- cells are clonal, these cells were reconstituted with GFP-tagged FAK to establish causality with FAK-/- phenotypes (Fig. 5F). Immunoblotting of cell lysates showed that active β-catenin (not phosphorylated at Ser33/37/Thr41) and Myc levels were restored by GFP-FAK re-expression in OVCAR3 FAK-/- tumorspheres. Reduced β-catenin and Myc within OVCAR3 FAK-/- AB21 cells is consistent with the sensitivity of these cells to CP-, JQ1-, and MLN8237-induced cytotoxicity. GFP-FAK promoted tumorsphere formation, rescued ALDH activity, and restored ALDH-1A2 and ALDH-1B1 protein levels in OVCAR3 FAK-/- AB21 cells (Figs. 5G-J). FAK knockout resulted in an approximate 10-fold reduction in OVCAR3 tumorsphere EC50 to CP (Fig. 5K). Notably, GFP-FAK expression rescued FAK-/- sensitivity to CP. These results show that FAK promotes OVCAR3 CSC phenotypes and CP resistance.

### KMF FAK inactivation and reconstitution link intrinsic FAK activity to β-catenin

Since KMF tumorspheres exhibit elevated β-catenin activity and CSC phenotypes (Fig. 1), CRISPR/Cas9 targeting was next used to inactivate the murine *PTK2* exon 4 in KMF cells. Sanger sequencing confirmed unique deletions/ insertions predicted to terminate FAK protein translation in each of four *PTK2* alleles identified in KMF cells (Supplemental Fig. S16). Exome sequencing of FAK-/- clone KT13 (90% of exons sequenced at 100X) detected only 165 unique variants, including the four *PTK2* alterations, indicating that CRISPR targeting was specific and that the KT13 genome is similar to KMF cells (Supplemental Table S8). CRISPR editing eliminated expression of FAK, but not the closely related kinase Pyk2 (Fig. 6A). FAK-/- KT13 cells exhibited sensitivity to CP-, MLN8237-, or MSAB-induced cytotoxicity (Fig. 6B) and displayed altered total β-catenin protein levels compared to actin (Fig. 6A). Together, our studies support a connection between FAK, β-catenin, and Myc in both OVCAR3 and KMF cells.

**Figure 6.**
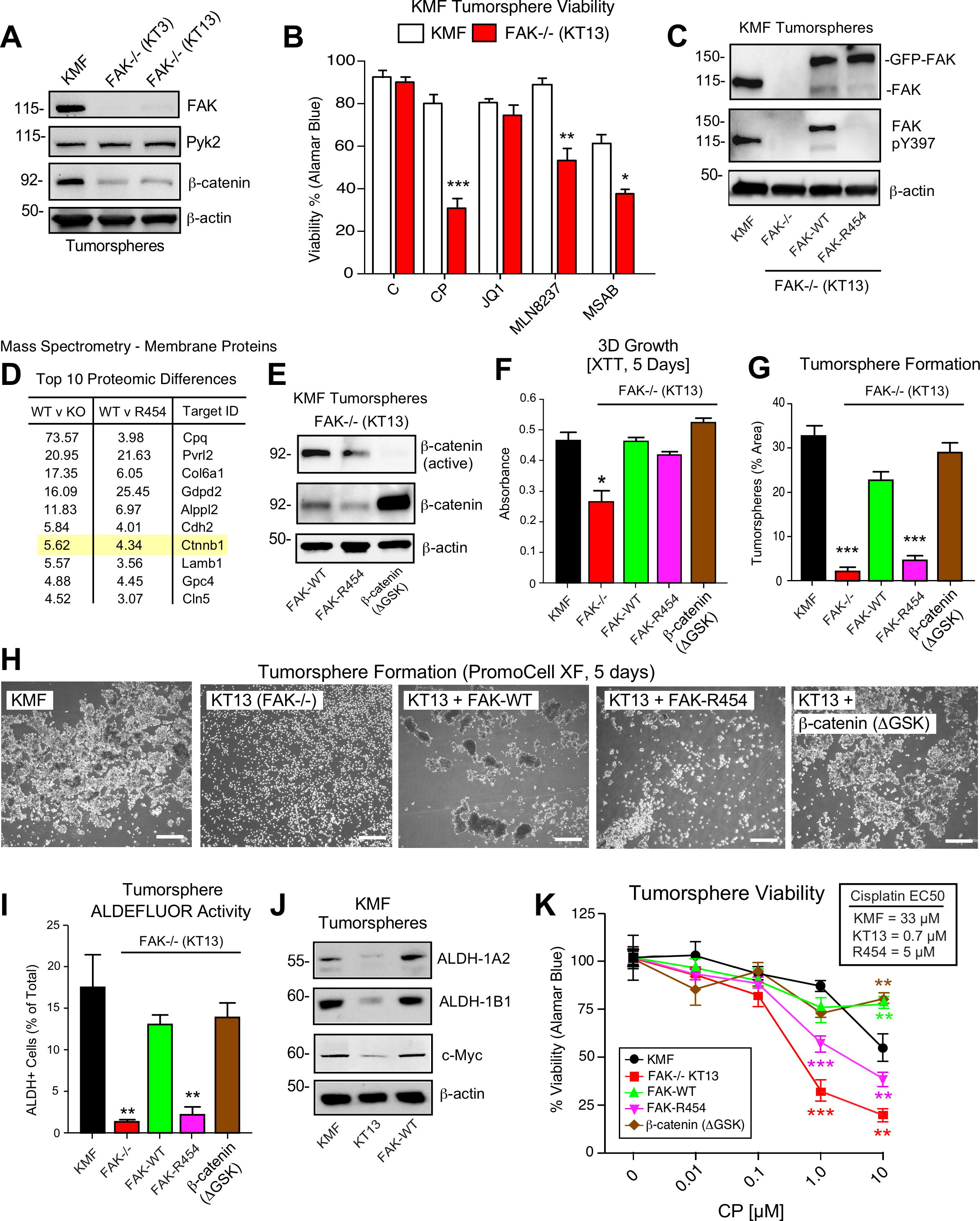
KMF FAK knockout and re-expression link intrinsic FAK activity and β-catenin to CSC phenotypes and CP resistance. **(A)** Immunoblotting of KMF, KT3 and KT13 cell lysates for FAK, Pyk2, β-catenin, and actin. **(B)** KMF and FAK-/- (clone KT13) cell viability treated with DMSO (control), CP (1 μM), JQ1 (1 μM), MLN8237 (1 μM), or MSAB (10 μM) after 72 h as measured by Alamar Blue. Values are means (± SEM, * P<0.05, ** P <0.01, *** P <0.001, one-way ANOVA with Fisher’s LSD multiple comparison test) for 3 independent experiments. **(C)** Immunoblotting for pY397 FAK, FAK, and actin in lysates of KMF, FAK-/- KT13, GFP-FAK-WT, and GFP-FAK R454 re-expressing cells. **(D)** Top ten proteomic differences (fold-change) detected by mass spectroscopy of membrane associated proteins in FAK-/- KT13, GFP-FAK-WT, and GFP-FAK R454 re-expressing cells. **(E)** Immunoblotting for active β-catenin, β-catenin, and actin in lysates of FAK-/- KT13 cells stably expressing GFP-FAK WT, GFP-FAK R454, or stabilized β-catenin (ΔGSK). **(F)** Growth of the indicated KMF, FAK-/- KT13, or the indicated reconstituted cells in PromoCell XF for 5 days. Values are means (± SEM, * P <0.05, one-way ANOVA) from 2 independent experiments. **(G)** Tumorsphere formation in PromoCell XF (5 days). **(H)** Representative images of the indicated cells in tumorsphere formation assays. Scale is 1 mm. **(I)** ALDEFLUOR activity. Values are means (± SEM **P<0.01, *** P <0.001, one-way ANOVA with a Tukey’s multiple comparisons test) of 3 (panel G) or 4 (panel I) independent experiments. **(J)** Immunoblotting for ALDH-1A2, ALDH-1B1, Myc, and actin in the indicated cell lysates. **(K)** Viability (Alamar Blue) of KMF (black circles), KT13 FAK-/- (red squares), GFP-FAK WT (green triangle), GFP-FAK R454 (magenta triangle), and ΔGSK β-catenin (brown diamond) expressing cells treated with increasing CP concentrations for 5 days. Values are means (± SEM, **, P <0.01, *** P <0.001, two-way ANOVA with a Bonferroni’s multiple comparisons test) from 3 independent experiments. EC50 values were determined independently and calculated using Prism (Graphpad).

As pharmacological FAK inhibition blocked KMF CSC phenotypes *in vitro* and *in vivo* (Figs. 3 and 4), a GFP-FAK point mutant (R454) eliminating FAK kinase activity (Sulzmaier et al., 2014) or GFP-FAK wildtype (WT) were stably-expressed in KMF FAK-/- KT13 cells (Fig. 6C). In 3D anchorage-independent conditions, GFP-FAK WT but not GFP-FAK R454 was phosphorylated at Y397 (Fig. 6C), thus implicating intrinsic kinase activity in FAK Y397 phosphorylation. Proteomic mass spectrometry analyses revealed elevated levels of extracellular matrix (*Col6a1* and *Lamb1*), surface receptors (*Pvrl2* and *Cdh2*), and β-catenin (*Ctnnb1*) in FAK-WT compared to FAK-/- (KO) and FAK-R454 cells (Fig. 6D and Supplemental Table S9). These results support the importance of intrinsic FAK activity in maintaining β-catenin protein levels in KMF cells.

To test whether enhanced β-catenin signaling could complement FAK-/- KMF cell phenotypes, an activated β-catenin point-mutant (ΔGSK) lacking the regulatory GSK3βphosphorylation sites (Barth, Stewart et al., 1999) was stably expressed in FAK-/- KT13 cells (Fig. 6E). β-catenin ΔGSK was over-expressed but not detected by antibodies to active β-catenin, likely due to disruption of the antibody epitope site (ΔGSK). FAK-/- KT13 cells did not form colonies in methylcellulose and exhibited decreased growth in 3D conditions (Fig. 6F). However, KT13 cells expressing GFP FAK-WT, GFP FAK-R454, or β-catenin ΔGSK grew equivalently in tumorsphere media to parental KMF cells (Fig. 6F). Notably, FAK R454 cells grow in 3D culture whereas parental KMF cells treated with FAKi exhibit growth defects (Fig. 3 and Supplemental Fig. S6).

Nevertheless, KT13 3D tumorsphere formation was significantly enhanced by FAK-WT and β-catenin ΔGSK but not FAK R454 re-expression (Figs. 6G and H). FAK-WT restored total ALDEFLUOR activity, ALDH-1A2, ALDH-1B1, and Myc protein levels in FAK-/- KT13 cells equivalent to parental KMF cells (Figs. 6I and J). Expression of FAK-WT and β-catenin ΔGSK but not FAK R454 significantly enhanced FAK-/- KT13 CP resistance *in vitro* (Fig. 6K). Together, these results link intrinsic FAK activity and β-catenin in supporting KMF CSC and intrinsic CP resistance phenotypes *in vitro*.

### Intrinsic FAK activity promotes KMF tumor growth

Although FAKi decreased tumor growth and inhibited secondary KMF tumor formation in mice (Fig. 4), it remained unclear whether *in vivo* FAKi effects were mediated via inhibition of FAK activity in tumor, stroma, or multiple cell types. Parental KMF, FAK-/- KT13, and KT13 FAK-WT cells were labeled with a dual reporter (luciferase and dTomato) and injected within the intraperitoneal cavity of C57Bl6 mice to test whether FAK is essential for KMF tumor formation (Fig. 7A). At Day 24, luciferase imaging revealed significant KMF tumor burden whereas FAK-/- KT13 tumors were only weakly detected (Fig. 7A and Supplemental Fig. S17). Direct enumeration of dTomato-positive cells isolated from a peritoneal wash at Day 28 revealed significantly fewer FAK-/- KT13 compared to mice bearing KMF and FAK-WT tumors (Fig. 7B). In an independent tumor experiment over 21 days, expression of FAK R454 did not promote FAK-/- KT13 tumor growth (Fig. 7C), supporting the essential importance of FAK activity for tumor growth *in vivo*. Surprisingly, β-catenin ΔGSK also did not promote FAK-/- KT13 tumor growth (Fig. 7C). This result contrasted with the rescue of FAK-/- KT13 tumorsphere formation, ALDEFLUOR activity, and CP resistance *in vitro* by β-catenin ΔGSK expression (Fig. 6). As tumor growth is a complex process, it is likely that β-catenin activation is one of several pathways downstream of FAK necessary for tumor growth.

**Figure 7.**
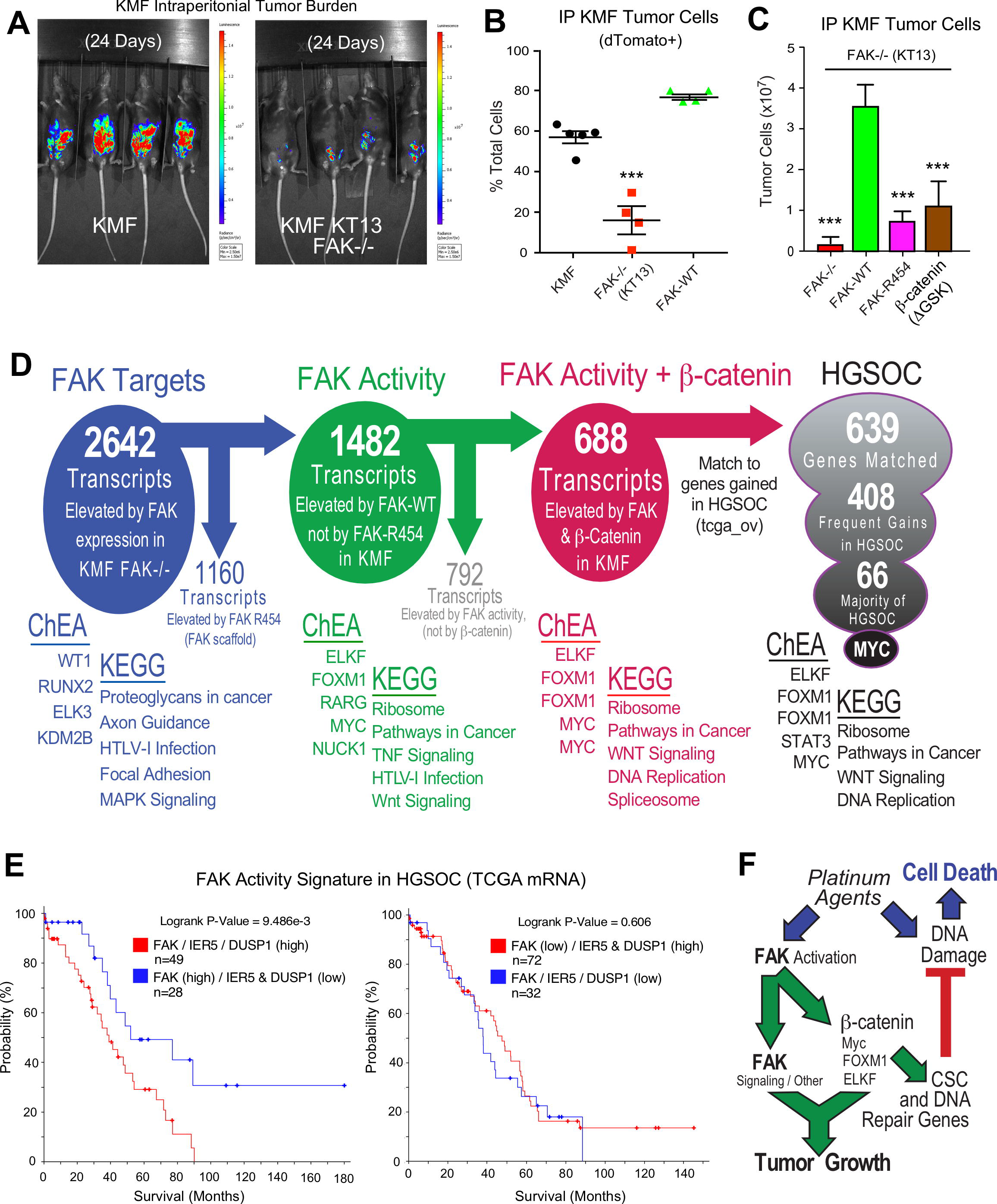
FAK activity and β-catenin promote a common Myc associated gene signature elevated in HGSOC. **(A)** IVIS imaging of C57Bl6 mice with dTomato+ and luciferase-expressing KMF or KT13 FAK-/- cells at experimental Day 24. (**B**) Flow cytometry analyses of peritoneal wash collected dTomato+ cells at Day 28 of mice bearing KMF (black), KT13 FAK-/- (red), and KT13 FAK-/- re-expressing FAK WT (green) cells. Values are means expressed as percent of total cells in peritoneal wash (± SEM, *** P<0.001, one-way ANOVA). (**C**) Intraperitoneal (IP) tumor growth of KT13 FAK-/- (red), GFP-FAK WT (green), GFP-FAK R454 (magenta), or ΔGSK β-catenin (brown) expressing cells. Values are means of CD45-negative tumor cells determined by flow cytometry (± SD, *** P<0.001, one-way ANOVA). **(D)** Summary of KMF RNA sequencing and filtering of differential gene expression. 2642 mRNAs were elevated (greater than log2) in FAK-WT versus KT13 FAK-/- cells. 1160 mRNAs were elevated in FAK R454 versus KT13 FAK-/- cells. This represents FAK scaffold or activity-independent group (blue). By subtraction of FAK R454 from FAK-WT targets, 1482 FAK activity-dependent targets were identified (green). 3899 mRNAs were elevated in ΔGSK β-catenin KT13 FAK-/- cells and by filtering against FAK activity mRNAs, 688 common FAK activity and β-catenin enhanced mRNA targets were identified (red). 639 of 688 murine KMF targets were matched to genes elevated in HGSOC. 408 targets were elevated in 20% of HGSOC patients and 66 targets were elevated in more than 50% of HGSOC patients. MYC and FAK show the highest frequency of genetic gain. Representative transcription factors identified by ChIP Enrichment Analysis (ChEA) and the top 5 Kyoto Encyclopedia of Genes and Genomes (KEGG) pathway enrichments are listed for filtered groups. **(E)** Kaplan-Meier analysis of the impact of expression of DUSP1 and IER5 (z>0.1) on overall patient survival expressing elevated (z>3) or lesser levels of FAK mRNA. (**F**) Signaling summary of death-inducing and paradoxical survival-sustaining FAK activation by platinum chemotherapy. FAK signaling to β-catenin, MYC, FOXM1, and KLF1 transcription factors support elevated mRNA levels for CSC and DNA repair genes hypothesized to support cellular resistance platinum DNA damage. Tumor cell intrinsic FAK kinase activity is essential for KMF tumor growth via signaling in a context-dependent manner.

### Transcriptomic analysis of common FAK activity-dependent and β-catenin mRNA targets

FAK controls various gene transcriptional networks (Serrels, McGivern et al., 2017, Sulzmaier et al., 2014). As FAK-/- KT13 KMF cells are deficient in a number of different phenotypes, we performed RNA sequencing from cells grown as 3D tumorspheres to determine potential FAK activity-dependent, -independent, and β-catenin-specific patterns of differential gene expression. Analyses of FAK-/- KT13 and FAK-WT re-expressing cells revealed 2642 mRNA transcripts increased two-fold or more by FAK and significant after multiple testing correction (Fig. 7D and Supplemental Table S10). By filtering out transcripts that were elevated in FAK R454 cells, 1482 genes were identified as FAK activity-dependent and showed KEGG pathway enrichment for *Ribosome, Pathways in Cancer, Tumor Necrosis Factor*, and *Wnt Signaling*. (Supplemental Table S11). Notably, MYC, FOXM1, and ELKF were identified as transcription factors involved in FAK activity-dependent target regulation by ChIP Enrichment Analysis (ChEA) (Supplemental Table S12).

RNA sequencing identified 3899 mRNA transcripts that were elevated 2-fold or more and significant after multiple testing correction by β-catenin ΔGSK expression in FAK-/- KT13 cells (Supplemental Table S10). Unbiased filtering of the 1482 FAK activity-dependent transcripts with those elevated by β-catenin ΔGSK revealed 688 transcripts as common targets regulated by FAK activity and β-catenin (Fig. 7D and Supplemental Table S10). ChEA analyses showed enhancement of ELKF, MYC and FOXM1 transcription factors with *Ribosome*, *Pathways in Cancer*, and *Wnt Signaling* as commonly enriched KEGG pathways. Several known targets that promote platinum resistance or stemness were identified as being regulated by FAK and β-catenin signaling in KMF cells (Table 2). These include *AMIGO2, CD44, DUSP1, FN1, and IER5* associated with platinum resistance and *EIF4BP1, GAB1, LIMA1, and RCL1* associated with stemness, respectively.

**Table 2.**
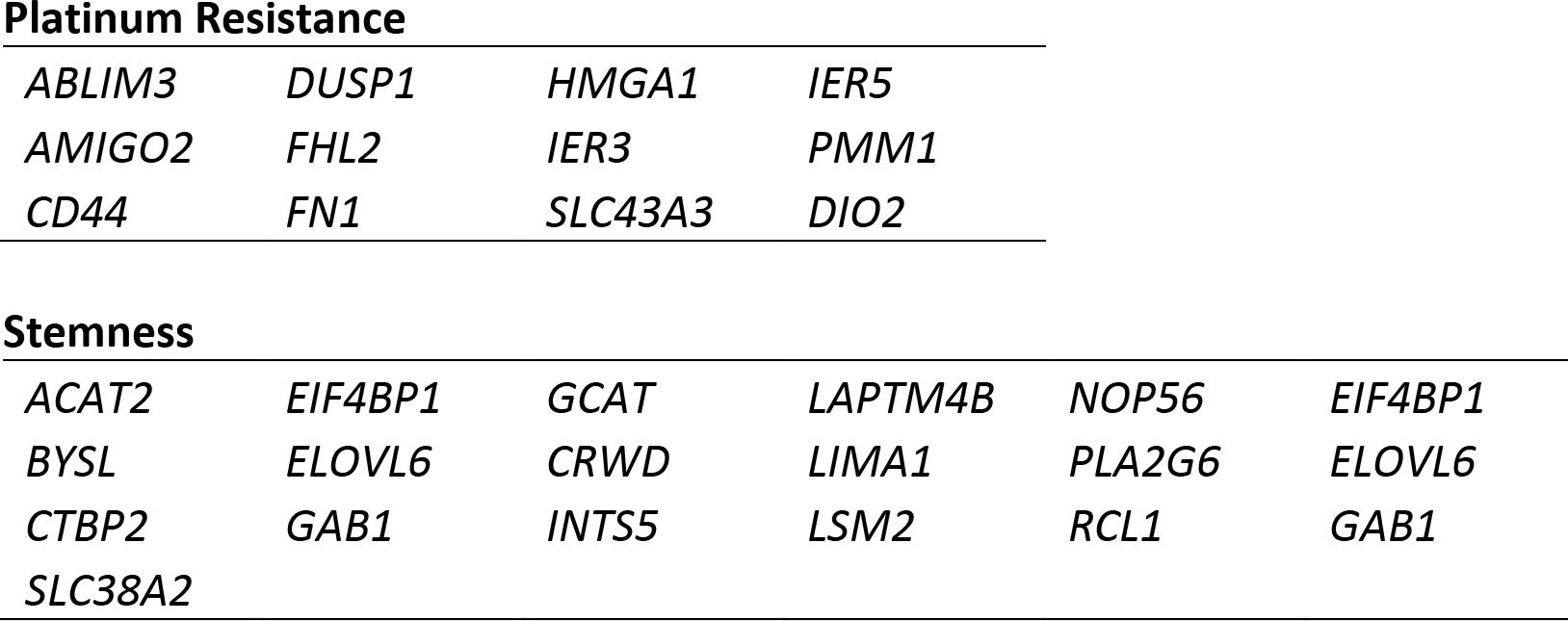
List of FAK activity-induced mRNA targets in KMF cells identified by GSEA analyses.

|  |  |  |  |
| --- | --- | --- | --- |
| *ABLIM3* | *DUSP1* | *HMGA1* | *IER5* |
| *AMIGO2* | *FHL2* | *IER3* | *PMM1* |
| *CD44* | *FN1* | *SLC43A3* | *DIO2* |

**Stemness**
|  |  |  |  |  |  |
| --- | --- | --- | --- | --- | --- |
| *ACAT2* | *EIF4BP1* | *GCAT* | *LAPTM4B* | *NOP56* | *EIF4BP1* |
| *BYSL* | *ELOVL6* | *CRWD* | *LIMA1* | *PLA2G6* | *ELOVL6* |
| *CTBP2* | *GAB1* | *INTS5* | *LSM2* | *RCL1* | *GAB1* |
| *SLC38A2* |  |  |  |  |  |

### Connections between FAK activity and MYC in HGSOC

To explore potential similarities between murine KMF cells and HGSOC, the 688 common FAK activity- and β-catenin-enhanced mRNA targets were filtered against genes elevated in HGSOC. Notably 639 targets (93%) were matched, 408 genes (61%) were elevated in at least 20% of TCGA tumor samples, and 66 genes were elevated-amplified in greater than 50% of patients (Supplemental Table S13). MYC was the most common gene target (80%) with FOXM1 and ELKF transcription factor targets enriched by ChEA analyses. KEGG enrichment analyses identified *Ribosome, Pathways in Cancer, Wnt Signaling*, and *DNA Replication* pathways. While many of these 66 genes were prognostic for poor outcome independent of FAK, the expression of immediate early response 5 (IER5) and dual specificity phosphatase 1 (DUSP1) were selectively associated with poor outcome among patients with elevated FAK expression (Fig. 7E) (logrank P = 9.486e-3). Although a role for IER5 in ovarian cancer is not yet described, elevated DUSP1 has been linked to platinum resistance (Kang, Nagaraja et al., 2016). Taken together, our transcriptomic and cell biological analyses of KMF cells have identified a FAK activity-dependent, β-catenin and Myc associated gene signature supporting CSC and platinum resistance signaling pathways in ovarian cancer. KMF cells are a unique murine model with profound similarities to HGSOC and will be made available to the research community.

## Discussion

Patients with platinum resistant ovarian cancer have minimally effective treatment options, complex tumor genomes, and few targetable oncogenic mutations (Patch et al., 2015). Gene breakage, gains, or losses are common drivers of tumor cell phenotypes. We provide evidence from a new in vivo-evolved murine ovarian cancer model termed KMF - denoting gains in <u>K</u>-Ras, <u>M</u>yc, and <u>F</u>AK genes - for the functional significance of *PTK2* (FAK) gains in ovarian cancer cell and tumor biology. KMF cells are notable for aggressive tumor growth, tumorsphere formation *in vitro*, elevated FAK Y397 phosphorylation, increased β-catenin and ALDH activities, deficiencies in p53 regulation, and increased resistance to CP cytotoxicity compared to parental ID8 cells. In both KMF and human OVCAR3 ovarian carcinoma cells, we identify tumorsphere associated non-canonical FAK signaling as supporting CSC phenotypes and intrinsic CP resistance.

Although *MYC* amplification at 8q24.21 is associated with a poor prognosis (GoodeChenevix-Trench et al., 2010), less is known about *PTK2* amplification at 8q24.3. We show that over 70% of HGSOC patient tumors contain gains at both *PTK2* and *MYC* loci, that *PTK2* copy number parallels FAK mRNA and protein increases, and that elevated FAK mRNA levels are associated with decreased patient disease-free survival. We have identified a set of 36 genes associated with *PTK2* gain and increased frequency of HGSOC relapse. Additionally, we identify Myc as part of a set of 66 genes increased in KMF cells in a FAK kinase-dependent manner that are also elevated in the majority of HGSOC patients. Notably, in this 66 gene set, elevated *PTK2*, *IER5*, and *DUSP1* expression as a group is associated with decreased HGSOC patient overall survival.

Another unexpected finding of our studies is that while platinum and taxane chemotherapy kills most ovarian tumor cells, we paradoxically find that FAK activation can occur in residual tumors of patients undergoing chemotherapy, in mouse tumors, and in isolated ovarian carcinoma tumorspheres after cisplatin chemotherapy (Fig. 7F). Consistent with studies linking chemotherapy to increased CSC survival (Wiechert et al., 2016), we find that FAK inhibition compromises ALDH activity and CSC generation *in vitro* and *in vivo.* Moreover, CP resistant cells can acquire FAK dependence for growth.

Our collective approaches, including pharmacological inhibition, FAK knockout, FAK re-expression, and complementation analyses show that FAK signaling sustains both intrinsic and acquired resistance to cisplatin chemotherapy in part via β-catenin activation and the elevation of MYC, FOXM1, and ELKF transcription factors supporting Wnt signaling and DNA replication genes involved in CSC survival and platinum resistance (Fig. 7F). Interestingly, a FAK to β-catenin signaling linkage has been identified as an adaptive chemotherapy resistance pathway in *BRAF* mutated colon cancer (Chen, Gao et al., 2018, Taylor & Schlaepfer, 2018). Although stabilized β-catenin ΔGSK expression in KMF FAK-/- cells activated a number of canonical Wnt target genes, this was unexpectedly insufficient to rescue KMF FAK-/- growth as tumors. Instead, we show that FAK expression and intrinsic activity are essential for KMF tumor growth and that elevated FAK activity and Y397 phosphorylation is an acquired and targetable cellular adaptation of cisplatin resistance in HGSOC.

Mechanistically, FAK functions as a signaling scaffold that is recruited to different cellular compartments in response to various stimuli (Sulzmaier et al., 2014). It has both kinase-dependent and -independent functions in regulating gene expression (Lim et al., 2008, Serrels et al., 2017). It is possible that β-catenin ΔGSK signaling requires FAK scaffold to support tumor growth *in vivo*, or that this stabilized β-catenin construct only functions as a weak oncogene (Barth et al., 1999). In cell culture, CP resistant cells acquired dependence on FAK activity to maintain proliferation as 3D tumorspheres without alterations in 2D growth. In contrast to other studies (Kolev et al., 2017), single agent pharmacological FAK inhibition did not promote apoptosis of ovarian CP resistant cells. Rather, the combination of FAK inhibition (genomic and pharmacological) with CP triggered apoptosis of CP resistant cells as tumorspheres *in vitro* and prevented CP-resistant tumor growth in mice. To this end, a clinical trial for recurrent CP resistant ovarian cancer termed ROCKIF (<u>R</u>e-sensitization of platinum-resistant <u>O</u>varian <u>C</u>ancer by <u>K</u>inase Inhibition of <u>F</u>AK, NCT03287271) will test whether the small molecule FAK inhibitor defactinib, in combination with carboplatin and paclitaxel chemotherapy, can provide benefit for this difficult to treat patient population.

## Materials and Methods

### Antibodies, Plasmids, and Reagents

Monoclonal (clone 4.47) and polyclonal (06-543) antibodies (EMD Millipore) were used to detect FAK. Rabbit monoclonal antibodies were used to detect FAK pY397 (Thermo clone 141-9, clone 31H5L17, and Abcam clone EP2160Y). Mouse FAK pY397 peptide (Abcam AB40145) was used for blocking. Monoclonal antibodies were used to detect E-cadherin (Cell Signaling Technologies [CST] clone 4A2), β-actin (Sigma clone AC-74 or Proteintech clone 7D2C10), β-catenin (CST clone D10A8), non-phospho (active) β-catenin (Ser33/Ser37/Thr41) (CST clone D13A1), Myc (CST clone D84C12), Pyk2 (CST clone 5E2), p21CIP1 (Santa Cruz Biotechnology [SCB}, clone F5), GFP (SCB clone B2), p53 (SCB Pab 240), and α-tubulin (Sigma clone DM1A). Polyclonal antibodies were used to detect ALDH-1A1 (Abcam ab23375), ALDH-1A2 (ProteinTech 13951), ALDH-1B1 (ProteinTech 15560), ALDH-3B1 (ProteinTech 19446), Ki67 (Abcam ab15580), PAX8 (Proteintech 10336-1-AP), p53 (Abcam ab26), β-catenin pY142 (Abcam ab27798), and cyclin D1 (CST 2922).

The dTomato with luciferase lentiviral vector, pUltra-Chili-Luc, was a gift from Malcolm Moore (Addgene #48688). The lentiviral vector MSCV-β-catenin (ΔGSK-KT3)-IRES-GFP was a gift from Tannishtha Reya (Addgene #14717). The CRISPR/Cas9 plasmid pSpCas9n(BB)-2A-Puro was a gift from Feng Zhang (Addgene #48141). Lentiviral vectors for GFP-FAK wildtype and FAK kinase-dead (R454) in pCDH-CMV-MCS-Puro (System Biosciences) were used as described (Chen et al., 2012). FAKi VS-4718 and VS-6063 (defactinib) were from Verastem Inc. FAKi, cisplatin (Enzo Life Sciences), JQ1 (MedChemExpress), MLN8237 (Selleckchem), MSAB (Sigma), wortmannin (Sigma), U0126 (Sigma) or staurosporine (Calbiochem) were dissolved in DMSO for *in vitro* studies. VS-4718 was suspended in 0.5% carboxymethyl cellulose (Sigma) and 0.1% Tween 80 (Sigma) in sterile water and administered twice daily by oral gavage for tumor experiments. Cisplatin and paclitaxel (APP Pharmaceuticals) were from the UCSD Moores Cancer Center Pharmacy.

### Cells

Human ovarian carcinoma A2780, A2780-CP70, and OVCAR10 cell lines were from Denise Connolly (Fox Chase Cancer Center, PA). NIH OVCAR3 cells were from the Division of Cancer Treatment & Diagnosis Tumor Repository, National Cancer Institute (Frederick, MD), murine ovarian ID8 cells were from Katherine Roby (University of Kansas Medical Center), and KMF cells were isolated from peritoneal ascites of ID8-injected C57Bl6 tumor-bearing mice as described (Ward et al., 2013). Intermittent CP exposure (10 μM for 24 h), cell recovery (7 days), and repeated exposure-recovery (5 times) was used to generate OVCAR10-CP cells.

OVCAR3 FAK knockout cells were generated using CRISPR/Cas9 targeting. pSpCas9n(BB)-2A-Puro was used to deliver guide RNAs (ACTGGTATGGAACGTTCTCC and TGAGTCTTAGTACTCGAATT) targeting exon 3 of human *PTK2*. Transfected cells were enriched by puromycin (1 μg/ml, 3 days) and clones selected by dilution. Loss of FAK expression was verified by immunoblotting. DNA sequencing was used to verify insertions/deletions introducing stop codons in *PTK2* exon 3. FAK re-expressing cells were generated by lentiviral transduction of OVCAR3 FAK KO clone AB21, puromycin selection, enrichment by flow cytometry, and GFP-FAK protein expression verified by immunoblotting. KMF FAK knockout cells were generated by CRISPR/Cas9 targeting. pSpCas9n(BB)-2A-Puro was used to deliver two independent guide RNAs (ACTTACATGGTAGCTCGCGG and CACTCCCACAGCCATCCTAT) targeting exon 4 of murine *PTK2*. Transfected cells were enriched by puromycin selection (3.5 μg/mL for 24 h) and clones selected by dilution. Loss of FAK expression was verified by immunoblotting. DNA sequencing was used to verify insertions/deletions introducing stop codons in murine *PTK2* exon 4. GFP-FAK-WT, GFP-FAK-R454, and ΔGSK β-catenin (GFP expressed independently) were generated by lentiviral (for FAK) or retroviral (for β-catenin ΔGSK) transduction, puromycin or hygromycin selection, enrichment by flow cytometry, and protein expression verified by immunoblotting. For adherent 2D growth, cells were maintained in DMEM (OVCAR10, OVCAR10-CP, ID8, and KMF) or RPMI 1640 (A2780, A2780-CP70, and OVCAR3) supplemented with 10% fetal bovine serum (FBS, Omega Scientific), 1 mM non-essential amino acids, 100 U/ml penicillin, and 100 μg/ml streptomycin on cell culture-treated plastic plates (Costar). For growth as tumorspheres, cells were seeded in poly-hydroxyethyl methacrylic acid (poly-HEMA) coated Costar plates (non-adherent) in serum-free CSC media (3D Tumorsphere Medium XF, PromoCell GmbH) at dilutions recommended by the manufacturer. Prior to tumor initiation experiments, KMF, A2780, and A2780-CP70 cells were transduced with a lentiviral vector expressing dTomato and luciferase (pUltra-Chili-Luc) or mCherry (pCDH-CMV-MSCI) and enriched by fluorescence sorting.

### 2D and 3D Cell Growth Assays

For 2D growth, cells were seeded (3×10^5^ cells per well) in tissue culture-treated 6-well plates (Costar). At the indicated time, cells were enumerated and stained with Trypan blue (ViCell XR, Beckman). Alternatively, cell metabolic activity was measured by a colorimetric XTT assay (Sigma). For 3D tumorspheres, cells were seeded at 10,000 cells/ml equivalent in poly-HEMA-coated 6-, 24-, or 96-well plates (Costar) for 5 days. At the indicated times, 3D tumorspheres were phase-contrast imaged (Olympus CKX41), enumerated (ViCell XR), or collected by centrifugation. Spheroid size was determined using Image J (NIH). Alternatively, cell metabolic activity was measured by a colorimetric XTT assay (Sigma). For methylcellulose colony formation, cells were suspended in 1% methylcellulose diluted in 2D growth media, 10^4^ plated in 6-well poly-HEMA-coated plates, and colony formation analyzed after 21 days. Cells from methylcellulose colonies were collected by dilution-dispersion in PBS, centrifugation at 400 xg, and washed in PBS prior to enumeration or cell lysis. Cells were used at passage 10 to 35 and mycoplasma testing was performed every 3 months.

### Patient Tumor Samples

De-identified human OC tissue specimens from consented patients were obtained from the Fox Chase Cancer Center (FCCC) Biosample Repository Facility (BRF) under Institutional Review Board (IRB) approved protocols (IRB 11-866 and IRB 08-851). FCCC staff queried the BRF sample database to identify participants that received carboplatin and paclitaxel neoadjuvant chemotherapy. Biopsy specimens were obtained from FCCC Surgical Pathology, sectioned, H&E stained, and reviewed by a board-certified pathologist. FFPE blocks from the biopsy and the corresponding surgical resection blocks banked by the BRF were cut to obtain one H&E stained slide and six additional unstained sections. One section each from pre-treatment biopsy and post-neoadjuvant treatment surgical resection specimen was stained for Pax8 by the FCCC Histopathology Facility. The remainder of unstained slides were sent to UCSD for additional staining performed under UCSD IRB-approved protocol (IRB 110805).

### Immunohistochemistry

Mouse tumors were divided into thirds and either processed for protein lysates, fixed in formalin, or frozen in optimal cutting temperature compound. For immunohistochemical staining, paraffin-embedded tumors were sectioned, mounted onto glass slides, deparaffinized, rehydrated, processed for antigen retrieval, and peroxidase quenched as described (Tancioni et al., 2014). Tissues were blocked (PBS with 1% BSA, and 0.1% Triton X-100) for 45 minutes at room temperature and incubated with anti-PAX8 (1:200), anti-FAK (1:200), anti-Ki67 (1:500), anti-active β-catenin (1:800) or anti-pY397 FAK (1:100) in blocking buffer overnight. FAK pY397 antibodies were pre-incubated with 200-fold molar excess of FAK pY397 peptide (Abcam) for 12 hours at RT prior to use in IHC staining. Processing with biotinylated goat-anti-rabbit or goat-anti-mouse IgG, Vectastain ABC Elite, and diaminobenzidine were used to visualize antibody binding. Slides were counterstained with hematoxylin. Colon or breast carcinoma tumor samples were used as controls for active β-catenin staining. High resolution digital scans were acquired (Aperio CS2 scanner) using Image Scope software (Leica Biosystems). Images were also acquired using an upright microscope (Olympus BX43) with a color camera (Olympus SC100). A board-certified pathologist evaluated H&E, Pax8, pY397 FAK, or Ki67 stained images of patient tumor samples in a blinded manner. Quantification was performed using Aperio Image Analysis software (v12.3.0.5056) using the positive pixel count (v9) algorithm. Pax8-positive regions were identified and then these regions were manually-transposed onto images from FAK pY397-stained serial section slides. Average intensity (I-avg) values were obtained and percent FAK pY397 was calculated.

Frozen tumors were thin sectioned (7 μm) using a cryostat (Leica), mounted onto glass slides, fixed with acetone (or with 4% paraformaldehyde) for 10 min, permeabilized (PBS with 0.1% Triton) for 1 minute, and blocked (PBS with 8% goat serum) for 2 hours at room temperature. Sections were incubated in anti-ALDH1A1 (1:100) or anti-pY397 FAK (1:100) in PBS with 2% goat serum overnight. Antibody binding was detected with goat anti-rabbit conjugated with Alexa Fluor-488. Cell nuclei were visualized using Hoechst 33342 stain (Thermo). Images were sequentially captured at 20X magnification (UPLFL objective, 1.3 NA; Olympus) using a monochrome charge-coupled camera (ORCA ER; Hamamatsu), an inverted microscope (IX81; Olympus), and Slidebook software (v5.0, Intelligent Imaging). Images were pseudo-colored, overlaid, merged using Photoshop (Adobe), and quantified using Image J (NIH).

### Statistics

Statistical difference between groups was determined using one-way or two-way ANOVA with Tukey, Bonferroni’s or Fisher’s LSD post hoc analysis. Differences between pairs of data were determined using an unpaired two-tailed Student’s t test. For the IHC analysis the differences between pairs of data were calculated using a paired two-tailed Student’s t test. All statistical analyses were performed using Prism (GraphPad Software, v7). *P*-values of <0.05 were considered significant.

## Abbreviations

3D: three-dimensional
ALDH: aldehyde dehydrogenase
ChEA: ChIP Enrichment Analysis
CP: cisplatin
CPT: cisplatin and paclitaxel
CRISPR: Clustered regularly interspaced short palindromic repeats
CSC: cancer stem cell
DEAB: diethylaminobenzaldehyde
FAK: focal adhesion kinase
FAKi: focal adhesion kinase inhibitor
HGSOC: high grade serous ovarian cancer
IHC: immunohistochemical
IP: intraperitoneal
KEGG: Kyoto Encyclopedia of Genes and Genomes
KO: knockout
poly-HEMA: poly-hyodroxyethyl methacrylic acid
pY397 FAK: FAK Y397 phosphorylation
TCGA: The Cancer Genome Atlas
TIF: tumor initiation frequency
Y: tyrosine

## Acknowledgements

We thank our colleagues for helpful discussion and comments. We thank R. Winters (FCCC BRF manager) and D. Flieder (FCCC Pathology) for identifying, reviewing the cases, and for procuring patient tumor samples used in this study. This work was supported by grants NIH RO1CA180769, NIH RO1CA102310, NIH RO1CA107263, NIH UL1TR001442, NIH P30CA023100 and from charitable donations from Nine Girls Ask. C. Diaz-Osterman and F.J. Sulzmaier were supported by NIH training grant (T32-CA121938). C. Jean received support from the Ligue Nationale Contre le Cancer (LNCC). L. Bean, K. Anderson, K. Taylor, and A. Barrie are fellows of the UCSD Reproductive Medicine Gynecologic Oncology program. K Taylor and A. Barrie are Gaines Gynecologic Oncology Fellows. E. Cordasco was supported in part by NIH T36 GM095349. B Győrffy was supported by the NVKP_16-1-2016-0037, 2018-1.3.1-VKE-2018-00032 and KH-129581 grants. D. Connolly was supported by NCI P30 CA006927, NCI CA195723, DOD W81XWH-16-1-0142 and charitable donations from The Roberta Dubrow Fund and The Main Line Chapter of the Board of Associates.

## Authors Contributions

Designing research studies: CDO, DO, EGK, KJT, AMB, LMB, FJS, CJ, IT, XLC, MO, DCC, AM, DGS, and DDS

Conducting experiments: CDO, DO, EGK, KJT, AMB, LMB, FJS, CJ, IT, KA, SU, EAC, JL, XLC, MO, PR, VNK, AM

Analyzing data: CDO, DO, EGK, KJT, AMB, SJ, LMB, FJS, CJ, IT, KA, SU, JL, XLC, GF, MO, PR, JH, AMM, GX, KMF, VNK, DTW, JAP, BG, MTM, DCC, AM, DGS, DDS

Providing reagents or expertise: GF, JH, DTW, JAP, and DCC.

Writing the manuscript: CDO, DGS, and DDS. Study supervision: MTM, DCC, AM, DGS, and DDS.

## The Paper Explained

### Problem

Ovarian cancer remains the leading cause of death from gynecologic malignancies. Although platinum-based chemotherapy is part of standard-of-care treatment and is effective at creating DNA adducts and triggering cell apoptosis, subpopulations of tumor cells can survive this stress. Three in four women with high grade serous ovarian cancer (HGSOC) will recur, develop platinum resistance, and succumb to disease. Approaches to re-sensitize these tumors to platinum chemotherapy, or prevent tumor escape, are critically needed.

The *PTK2* gene at 8q24.3, encoding focal adhesion kinase (FAK), is frequently amplified in ovarian tumors. FAK is a cytoplasmic tyrosine kinase promoting cell survival signaling. FAK is associated with HGSOC progression, but the determinants of FAK pathway dependence in tumors remain unknown.

### Results

We molecularly characterize a new murine model of ovarian cancer that displays spontaneous acquired similarities to HGSOC, including gains in the <u>*K*</u>-*Ras*, <u>*M*</u>yc, and <u>F</u>AK genes (KMF). Our collective approaches in human ovarian carcinoma and KMF cells, including pharmacological inhibition, FAK knockout, FAK re-expression, complementation, and bioinformatic analyses reveal that FAK signaling sustains both intrinsic and acquired resistance to platinum chemotherapy in part via β-catenin activation and the elevation of *MYC, FOXM1*, and *ELKF* (KLF1) transcription factors supporting Wnt signaling and DNA replication genes involved in cancer stem cell survival and platinum resistance. FAK is unexpectedly activated in KMF and HGSOC tumor cells surviving neoadjuvant chemotherapy and we find that this is an acquired and targetable cellular adaptation of platinum resistance. The combination of FAK inhibition (genomic & pharmacologic) with cisplatin triggered apoptosis of platinum-resistant cells as tumorspheres *in vitro* and in orthotopic models *in vivo*. Gene editing and rescue approaches confirmed that intrinsic FAK activity is essential for tumor growth in mice.

### Impact

We identified a set of 36 genes associated with *PTK2* gain and increased frequency of HGSOC relapse. Moreover, a set of 66 genes including Myc were elevated in a FAK kinase dependent manner correspond to genes gained in the majority of HGSOC patients. Within this gene set, increased *IER5* and *DUSP1* expression are associated with decreased survival of patients bearing 8q24.3 gains. A clinical trial for recurrent platinum resistant ovarian cancer termed ROCKIF (<u>R</u>e-sensitization of platinum-resistant <u>O</u>varian <u>C</u>ancer by <u>*K*</u>inase Inhibition of <u>F</u>AK, NCT03287271) will test whether the small molecule FAK inhibitor defactinib, in combination with carboplatin and paclitaxel chemotherapy, can provide benefit for this difficult to treat patient population.

## Data Availability

The Exome sequencing FASTA files have been deposited to the NCBI Gene Expression Omnibus under the accession numbers [in progress]. The RNA-Sequencing FASTQ files used in this paper have been deposited to the NCBI Gene Expression Omnibus under the accession numbers [in progress]. The mass spectrometry proteomics data have been deposited to the ProteomeXchange Consortium via the PRIDE (Perez-Riverol, Csordas et al., 2019) partner repository with the dataset identifier PXD013062.

## List of Supplemental Materials

Additional Materials and Methods

## Headings to Supplemental Figures (full legend on figure)

Fig. S1. List of ID8 and ID8-IP/KMF gene variants identified by exome sequencing.

Fig. S2. Analysis of MYC and PTK2 mRNA and protein expression as a function of genomic copy number.

Fig. S3. Deficient p53 regulation in KMF cells.

Fig. S4. Characterization of ID8 and KMF cells.

Fig. S5. FAK pY397 phosphorylation is maintained in Pax8-positive HGSOC tumors after neo-adjuvant chemotherapy.

Fig. S6. Small molecule FAK inhibition selectively prevents KMF 3D tumorsphere proliferation with effects on cell cycle but not cell apoptosis.

Fig. S7. Small molecule FAK inhibition selectively prevents OVCAR3 3D tumorsphere proliferation with effects on cell cycle but not cell apoptosis.

Fig. S8. FAK controls 3D tumorsphere cell cycle in paired cisplatin-resistant but not cisplatin-sensitive ovarian carcinoma cells.

Fig. S9. Dose-dependent inhibition of A2780-CP70 methylcellulose colony growth by VS-6063 FAK inhibitor (Defactinib).

Fig. S10. Elevated ALDEFLUOR activity in paired CP-resistant compared to CP-sensitive tumorspheres.

Fig. S11. Oral FAKi (VS-4718) administration decreases KMF tumor burden and KMF-associated ALDEFLUOR activity.

Fig. S12. Inhibition of A2780 tumor growth by cisplatin-paclitaxel (CPT) chemotherapy.

Fig. S13. Selective inhibition of ALDEFLUOR activity within A2780-CP70 peritoneal tumor cells in vivo by oral VS-4718 administration.

Fig. S14. Elevated FAK Y397 phosphorylation and ALDH staining in non-necrotic regions of CPT treated mice with A2780-CP70 tumors.

Fig. S15. Validation of methyl 3-(4-methylphenyl)sulfonyl]amino-benzoate (MSAB) induced inhibition of β-catenin.

Fig. S16. Sequencing validation of FAK CRISPR/Cas9-mediated knockout in murine KMF cells.

Fig. S17. Comparison of KMF, KT13, and KT13 + GFP-FAK WT orthotopic growth in C57Bl6 mice.

## Headings to Supplemental Tables

Table S1. ID8 unique unique variants listed and filtered.

Table S2. ID8-IP / KMF unique variants listed and filtered.

Table S3. ID8 and KMF shared variants filtered.

Table S4. KMF copy number alterations called.

Table S5. HGSOC PTK2 gene signature list.

Table S6. RNA sequencing - differentially expressed genes in ID8 and ID8-IP / KMF tumorspheres.

Table S7. Patient tumor samples pre- and post-neoadjuvant chemotherapy, qualitative IHC score, and summary of quantitative image analyses.

Table S8. KMF FAK-/- KT13 exome sequencing variants.

Table S9. Summary of mass spectrometry-detected proteomic changes in KMF KT13 FAK-/-, FAK-WT, and FAK R454 cells.

Table S10. RNA sequencing - differentially expressed genes in KMF, KT13 (FAK-/-), KT13 FAK-WT, KT13 FAK R454, and KT13 ΔGSK β-catenin tumorspheres.

Table S11. KMF KEGG pathway analyses.

Table S12. KMF ChEA transcription factor analyses.

Table S13. FAK activity associated targets in KMF cells matched to HGSOC.

Table S14. Primers used for quantitative real-time PCR.

