## Supplemental Methods for "FAK activity sustains intrinsic and acquired ovarian cancer resistance to platinum chemotherapy"

[1]Department of Obstetrics, Gynecology, and Reproductive Sciences, Moores UCSD Cancer Center, La Jolla, CA 92093
[2]Xiamen University, Life Sciences, Innovation Center for Cellular Stress Biology, China
[3]Department of Biochemistry, University of Turku, Finland
[4]Department of Medicine, UCSD Center for Computational Biology & Bioinformatics, La Jolla, CA 92093
[5]Verastem Incorporated, Needham, MA, 02494
[6]Institute of Enzymology, Hungarian Academy of Sciences, and Semmelweis University 2nd Dept. of Pediatrics, Budapest, Hungary
[7]Fox Chase Cancer Center, Philadelphia, PA 19111
[8]Department of Pathology, Moores UCSD Cancer Center, La Jolla, CA 92093
*authors contributed equally to this study

[9]Co-corresponding authors:

| | |
|---|---|
| David D. Schlaepfer, Ph.D. | Dwayne G. Stupack, Ph.D. |
| | |
| https://orcid.org/0000-0003-4814-9210 | https://orcid.org/0000-0003-4396-5745 |

Department of Obstetrics, Gynecology, and Reproductive Sciences
UCSD Moores Cancer Center
3855 Health Sciences Dr., MC 0803
La Jolla, CA 92093

**Cell Cycle Analysis**
Cells were collected as a single cell suspension by limited trypsin treatment, fixed in 70% ethanol and stored at −20°C overnight. Cells were incubated in 100 µl of PBS containing DNAse-free RNAse (100 µg/mL, Qiagen). After 45 minutes, propidium iodide (PI, 10 µg/mL) was added prior to flow cytometry and analyzed using FlowJo (v9.5.1) and ModFit LT (Verity Software House) software.

**β-catenin transcriptional activity**
293T cell transfection with a β-catenin DNA binding reporter (7X-TCF repeat sequence AGATCAAAGGgg) driving eGFP (7TGP, Addgene #24305, gift from Roel Nusse) was packaged with psPAX2 (gift from Didier Trono, Addgene #12260) and pMD2.G (gift from Didier Trono, Addgene #12259) using Fugene HD (Thermo) according to the manufacturer's instructions. Media was replaced after 6 h, conditioned media collected (36 h), centrifuged (500g), and target cells infected for 24 h with polybrene (8 µg/ml Sigma). Transduced cells were selected by puromycin (1 µg/ml, 5 days). Proliferating cells were seeded in 3D tumorsphere growth conditions (PromoCell XF), DMSO or GSK3β inhibitor (CHIR99021, Sigma, 3 µM) added after 24 h, and cells evaluated by flow cytometry (BD FACSCelesta) after 5 days.

**Viability and Cytotoxicity Studies**
For annexin V and 7-AAD staining, cells were cultured in adherent or non-adherent conditions as described above for 24 hours or 5 days, respectively, with increasing cisplatin treatment (0 to 100 µM). Cells were dissociated using limited trypsin treatment, washed by centrifugation, suspended in annexin-V binding buffer (10 mM Hepes, 140 mM NaCl and 2.5 mM $CaCl_2$) and incubated with allophycocyanin-conjugated Annexin V (eBioscience) and 7-aminoactinomycin D (7-AAD) for 10 minutes at RT prior to analysis on a FACSCalibur flow cytometer (BD Biosciences). Post-acquisition analyses were performed using CellQuest Pro (BD Biosciences) or FlowJo software (v9.5.1). For AlamarBlue (Life Technologies) assays, cells were cultured in 96-well poly-HEMA-coated plates as above followed by treatment with cisplatin, VS-4718, MSAB, JQ1, MLN8237 or cisplatin plus individual inhibitors for 5 days. AlamarBlue reagent (Life Technologies) was added to each sample (10% final concentration) and incubated at 37°C at 5% $CO_2$ for 24 hours. Viability was analyzed by resorufin production via absorbance at 570/600 nm using a Synergy HTX spectrophotometer (BioTek Instruments).

**Cisplatin Cytotoxicity**
Cells (10,000 in 90 µl) were plated in tissue culture-treated 96-well plates (Costar). At 24 hours, increasing concentrations of cisplatin was added in growth media (10 µl), and the number of viable cells determined at 72 hours using the CellTiter 96® AQueous One Solution Cell Proliferation Assay (Promega). Measurement of cell resistance to CP-induced cytotoxicity were performed by colorimetric XTT cell staining (Sigma). In 2D culture, OVCAR10-CP and OVCAR10 cells exhibit 10.9 ± 2.2 µM and 1.4 ± 0.6 µM EC50 values to CP treatment, respectively. A2780-CP70 and A2780 cells exhibit 62.2 ± 8.7 µM and 5.6 ± 3.2 µM EC50 values to CP treatment, respectively. EC50 values were calculated using Prism (v7, GraphPad).

**ALDEFLUOR Assay**
The ALDEFLUOR fluorescent reagent system (Stemcell Technologies) was used to measure cell-associated ALDH activity. Cells were cultured as tumorspheres, treated with the indication concentrations of cisplatin, VS-4718, MSAB, JQ1, MLN8237, wortmannin, or U0126 for 5 days, collected by centrifugation, dissociated by trypsinization, resuspended in Aldefluor assay buffer containing ALDH substrate (BODIPY-aminoacetaldehyde), and incubated for 45 minutes at 37°C

with or without the ALDH inhibitor diethylamino-benzaldehyde (DEAB). AldeRed substrate (EMD Millipore) was used with cells expressing GFP. Individual gates were used to determine the percentage of ALDEFLUOR-positive cells per experimental point relative to DEAB-inhibitor treated controls. For analysis of ALDH activity in ascites-associated cells, pooled isolates from peritoneal washings of each experimental group were dissociated by trypsinization, treated with red blood cell lysis buffer (Biolegend), and processed as described above.

**Quantitative RT-PCR**

Total RNAs were extracted using PureLink™ RNA Mini Kit (Thermo) and cDNA prepared using the High-Capacity cDNA Reverse Transcription Kit (Thermo) from 1 µg total RNA. Target transcripts were amplified using a LightCycler 480 (Roche Applied Science), Premix Ex Taq probe qPCR Kit, iTaq™ Universal SYBR® Green Supermix (Bio-rad) with cDNA template and primers according to manufacturer instructions. Target gene expression was normalized to 40S ribosomal protein S17 (RS17) as a housekeeping gene control. Transcript levels were calculated using the $\Delta\Delta$CT (cycle threshold) method.

**Protein Analyses**

Protein extracts of cells were prepared using a lysis buffer containing (25 mM HEPES, pH 7.5, 150 mM NaCl, 10% glycerol, 10 mM $MgCl_2$, 1 mM EDTA, 10 mM NaF, 1 mM $Na_3VO_4$) with 1% NP-40, 0.25% sodium deoxycholate, 0.1% SDS and protease inhibitors (Roche Diagnostics). Tumors were homogenized in lysis buffer without sodium deoxycholate using Precellys24 (Bertin Instruments) bead disruption. Total protein levels in lysates were determined through a bicinchoninic acid assay (Pierce), proteins were resolved by SDS-PAGE (NuPAGE 4-12 % Tris-Bis gels, Thermo), and transferred to polyvinylidene difluoride membranes (Immobilon-FL, Millipore) for immunoblotting. Levels of protein expression and/or phosphorylation were detected with specific primary antibodies and IRDye 680 goat anti-mouse and IRDye 800 goat anti-rabbit secondary antibodies. Protein bands were visualized and quantified using the Odyssey Infrared Imaging System (Li-Cor Biosciences). Alternatively, HRP-conjugated secondary antibodies were visualized by chemiluminescence detection (ChemiDoc, BioRad).

**Tumor Growth in Mice**

All animal experiments were performed in accordance with The Association for Assessment and Accreditation for Laboratory Animal Care guidelines and approved by the UCSD Institutional Animal Care and Use Committee (S07331). For analysis of KMF response to chemotherapy, 4 million pChili-luciferase-labeled KMF cells in 250 µl DMEM were suspended with 250 µl phenol red-free high concentration Matrigel (Corning, 354262) and intraperitoneal injected into 10-week-old female C57BL6 mice (Charles River). Tumor growth was monitored weekly via bioluminescent luciferase imaging (IVIS, Perkin Elmer). On Day 7, mice were randomized to a control (saline injection) or chemotherapy group receiving IP injection of cisplatin (3 mg/kg) plus paclitaxel (2 mg/kg) at Day 7, 14, and 21. On Day 42, omental-associated tumors were dissected, minced, dissociated by incubation with collagenase, and cultured in DMEM with 10% FBS containing 20 µg/ml ciprofloxacin. Omental-associated KMF cells were expanded-passaged three times, analyzed for growth resistance to cisplatin, and lysates probed for pY397 and total FAK levels by immunoblotting.

Orthotopic mCherry-labeled KMF tumor growth in the ovarian bursa of female C57Bl6 mice was performed as described (Ward, Tancioni et al., 2013). On Day 7, mice were randomized and administered vehicle or VS-4718 (50 mg/kg via oral gavage, BID). On Day 28, mice were euthanized, cells from a peritoneal wash were collected, tumor implants were visualized by fluorescent imaging, and solid tumors were dissected and stored in Aquix RS-I media (Aquix LTD) at 4°C overnight. Tumor tissue from vehicle or VS-4718-treated tumors were dissociated to single-cell suspension using GentleMax (Miltenyi). Cells were collected by centrifugation, filtered through a 100 µm nylon mesh, and resuspended in DMEM with 20% FBS for enumeration. Two hundred thousand live cells from each primary tumor were pooled together and aliquots with $10^5$, $1.4x10^4$, $2x10^2$ and 300 cells were prepared. Cells were washed with PBS and mixed with 100 µl PBS:Matrigel (1:1) and injected into the flanks of female immunodeficient hairless NOD SCID mice (Harlan). Tumor formation was assessed 7 weeks after implantation. The number of tumor-initiating cells was calculated using Extreme Limiting Dilution Analysis software (http://bioinf.wehi.edu.au/software/elda/).

A2780 or A2780-CP70 tumor growth was evaluated by IP injection of 4 million pChili-Luciferase-labeled cells mixed with Matrigel into 9-week-old female NOD SCID gamma mice (Jackson Laboratory). IVIS imaging (Day 4, 11, 18, and 23) was used to monitor tumor growth. On Day 5, mice were randomized to a control (saline injection); chemotherapy group (CPT) receiving IP injection of cisplatin (3 mg/kg) plus paclitaxel (2 mg/kg) at Day 5, 12, and 19; VS-4718 FAK inhibitor (100 mg/kg) via oral gavage twice daily (BID); or CPT plus FAK inhibitor treatment. At Day 24, mice were euthanized, omental tumors excised, and remaining peritoneal metastatic sites quantified by dTomato fluorescence using an OV100 Small Animal Imaging Station (Olympus) and ImageJ software.

For KMF intraperitoneal tumor growth, cells were transduced with a lentiviral vector expressing dTomato and luciferase (pUltra-Chili-Luc) and were enriched by FACS. Cells were mixed with PBS + 50% Matrigel (Growth factor reduced, Corning) for a final concentration of 5 x $10^6$ cells per 250 µL using 10 week old C57Bl6/N mice (Charles River). Tumor growth was monitored via luciferase bioluminescent imaging (IVIS, Perkin Elmer). At the indicated times, ascites-associated cells were recovered by peritoneal washings by injection and immediate removal of PBS (5 ml), followed by erythrocyte lysis (RBC lysis buffer, eBioscience), Accutase treatment (Corning), and total cell enumeration with trypan blue for viability (ViCell XR, Beckman). Flow cytometry was used to determine the percent dTomato+ and CD45- cells recovered as KMF tumor burden.

**Exome Sequencing and CNV analysis**

Exome sequencing was performed by Novogene (Beijing, China), using genomic DNA (1 µg) isolated from ID8 or KMF cells. Genomic DNA was sheared into 180-280 bp fragments using a Covaris Sonicator (Covaris). Exome enrichment and sequencing libraries were generated using Agilent SureSelect Mouse All Exon kit (Agilent Technologies) following manufacturer's recommendations. Each exome was sequenced using a 150 bp paired-end protocol on the Illumina HiSeq platform, generating 47M reads for the ID8 sample and 61M reads for the KMF sample. (https://software.broadinstitute.org/gatk/best-practices). Reads were aligned with BWA MEM 0.7.12 (Li & Durbin, 2009) to mouse genome GRCm38_68. Variants were called with GATK 3.4 according to the Broad Institute's best practices (https://software.broadinstitute.org/gatk/best-practices) (McKenna, Hanna et al., 2010). Processing after alignment was carried out with SAMtools v.1.1 (Li & Durbin, 2009). Variants were annotated with ANNOVAR (Wang, Li et al., 2010). Copy number variants were called from the same alignments with CNVkit (Talevich, Shain et al., 2016), visualized in the Integrative Genomics Viewer (Robinson, Thorvaldsdottir et al., 2011) using standard parameters, with ID8 as normal and KMF as tumor samples. Ninety percent of exons were sequenced at 100X.

**RNA Sequencing and Analyses**

Total RNA was isolated from cells growing in suspension using PureLink RNA Mini Kit (Thermo Fisher). Three independent samples of RNA were isolated from ID8 or KMF cells grown in 3D PromoCell XF media as tumorsperes for 5 days at various cell passages. RNA sequencing was performed by Novogene (Beijing, China). Three replicate RNA samples were obtained from KT13, KT13 + GFP-FAK WT, GFP-FAK R454 (kinase-inactive), and KT13 expressing a ΔGSK β-catenin. RNA library preparation was performed using NEB Next Ultra RNA Library Prep Kit (New England Biolabs). Each transcriptome was sequenced using a 150 bp paired-end protocol on the Illumina HiSeq platform. At least 60 million clean reads were generated per sample. Reads were mapped (>90%) to the reference genome using TopHat2 (Kim, Pertea et al., 2013). Novogene analyses used ClusterProfiler software for enrichment analysis, including GO Enrichment, DO Enrichment, KEGG and Reactome database Enrichment to analyze and visualize functional profiles of genomic coordinates, genes and gene clusters. Novogene performed differential expression analysis of two conditions/groups by using the DESeq2 R package. Only transcripts with FPKM >1 were used, with criterion being adjusted *P* value <0.05. Enriched datasets from the GFP-FAK WT and GFP-FAK 454 datasets were merged: common genes were sifted into a scaffolding dataset and unique genes were sifted into a kinase-dependent dataset.

Each dataset was subject to GSEA/MSigDB analysis. The Kinase-dependent transcripts were then merged with the genes elevated in the ΔGSK β-catenin set. The common set of elevated transcripts were grouped into a seperate FAK Kinase B-Cat dependent dataset, and GSEA analysis again performed. Finally, this dataset was compared with the total list of all genes gained in HGSOC by GISTIC (ov_tcga) for genes gained in at least one in 5 (408 gene dataset) or in the majority (63 gene dataset) of TCGA ovarian cancer patients. For each of the 66 genes, Kaplan - Meier overall survival was evaluated using the TCGA data set for based on expression of individual genes was evaluated with cbioportal.org using curated patient sample lists.

**Ovarian Cancer PTK2 Transcriptomic Survival Analysis**

Survival analysis was performed using a database of ovarian cancer samples (Penzvalto, Lanczky et al., 2014). The TCGA dataset was used to link copy number gains to gene expression (Cancer Genome Atlas Research, 2011). Samples with copy number gains were designated into one cohort and all remaining samples were designated into a second cohort. Gene expression was compared between cohorts using a non-parametric Mann-Whitney test. Genes with a fold change over two and a p-value below 1E-04 were accepted as statistically significant. The mean expression of all significant genes was computed for each sample and was used in subsequent analyses for the selected gene. Cox proportional hazards regression was performed for relapse-free survival and for overall survival. Correlation between mRNA expression and survival was assessed by using the Kaplan-Meier plotter (Gyorffy, Lanczky et al., 2012) for PTK2 mRNA levels in 1435 annotated ovarian cancer patient samples. Selections were: relapse-free survival, split patients by median, stage (all), histology (serous), grade (all), debulk (all), and chemotherapy treatments (all).

**Proteomics – Mass Spectrometry**

ECM-enriched protein extracts from tumorsphere cultures in PromoCell XF were prepared by trypsin digestion as described (Ojalill, Rappu et al., 2018). Peptides were separated by a nanoflow HPLC system (Easy-nLC1000, Thermo) coupled to a Q Exactive Hybrid Quadrupole-Orbitrap Mass Spectrometer (Thermo). A full MS (mass spectrum) scan over the mass-to-charge (m/z) range of 300-2000 with a resolution of 140,000 followed by data dependent acquisition of with an isolation window of 2.0 m/z and a dynamic exclusion time of 20 s was performed. The top 10 ions were fragmented by higher energy collisional dissociation (HCD) with a normalized collision energy of 27 and scanned over the m/z range of 200-2000 with a resolution of 17,500. After the MS2 scan for each of the top 10 ions had been obtained, a new full MS scan was acquired and the process repeated until the end of the 70-min run. Three repeated runs per sample were performed. Tandem mass spectra were searched using MaxQuant software (v1.5.2.8) against reviewed (SwissProt) mouse sequences of UniProtKB release 2018_08. Peptide-spectrum-match- and protein-level false discovery rates were set at 0.01. Carbamidomethyl (C), as a fixed modification, and oxidation (M, P, K) as dynamic modifications were included.

A maximum of two missed cleavages was allowed. The LC-MS profiles were aligned, and the identifications were transferred to non-sequenced or non-identified MS features in other LC-MS runs (matching between runs). The extracted ion intensities of all peptides matching to the same protein from the three technical replicates were summed together. The protein was determined as detected in the sample if its identification had been derived from at least two unique peptide identifications. The contaminant proteins (according to the contaminants listed in MaxQuant), reverse identifications, and identifications only by site were removed. Only the proteins containing “cell membrane”, “plasma membrane”, “cell surface”, “extracellular matrix” or “secreted” in the cellular component gene ontology or in the subcellular location definition in the UniProt database were included in the final list. Samples were normalized by sum of protein intensities in the final list.
