## Supplemental Table S7 for "FAK activity sustains intrinsic and acquired ovarian cancer resistance to platinum chemotherapy"

**Supplemental Table S7.** Patient tumor samples pre- and post-neoadjuvant chemotherapy, qualitative IHC score, and summary of quantitative image analyses.

|  | **Specimen ID** | **Tissue Type** | | **Neoadjuvant C-T (cycles)** | **Biopsy Collection Interval (months)** | **Qualitative PAX8+ (%)** | **Quantification (Average Intensity)** | | |
| --- | --- | --- | --- | --- | --- | --- | --- | --- | --- |
|  |  |  |  |  |  |  | ***FAK pY397+*** | ***PAX8+*** | ***Percent FAK pY397+ / Pax8+*** |
| 1 | 1013983  (pre) | Poorly differentiated adenocarcinoma | A |  |  | 85 | 136.1 | 114.1 | 119.3 |
|  | 1013983  (post) | High-grade carcinoma, favor serous type | T | 3 | 1 | 45 | 111.7 | 137.7 | 81.1 |
| 2 | 1014086  (pre) | Adenocarcinoma consistent with ovarian serous carcinoma | A |  |  | 60 | 135.1 | 117.6 | 114.9 |
|  | 1014086  (post) | High-grade Serous, carcinoma | T | 3 | 2 | 85 | 148.6 | 120.4 | 123.5 |
| 3 | 1014252  (pre) | Adenocarcinoma. | A |  |  | 50 | 131.1 | 120.7 | 108.7 |
|  | 1014252 (post) | High grade serous adenocarcinoma | T | 3 | 1 | 20 | 162.9 | 182.8 | 89.1 |
| 4 | 1016253  (pre) | Adenocarcinoma consistent with ovarian serous carcinoma | A |  |  | 35 | 120.7 | 104.6 | 115.5 |
|  | 1016253 (post) | Serous adenocarcinoma | T | 3 + Bev | 1 | 35 | 132.0 | 103.4 | 127.6 |
| 5 | 1016247  (pre) | Papillary serous carcinoma | A |  |  | 65 | 73.1 | 138.1 | 52.9 |
|  | 1016247  (post) | High-grade papillary serous carcinoma | T | 6 | 2 | 55 | 132.7 | 115.6 | 114.8 |
| 6 | 1013123  (pre) | High grade serous adenocarcinoma | T |  |  | 80 | 92.0 | 125.6 | 73.3 |
|  | 1013123- (post) | High grade serous adenocarcinoma | T | 3 + Bev | 2 | 80 | 104.7 | 119.5 | 87.6 |
| 7 | 1014109  (pre) | High grade serous adenocarcinoma | T |  |  | 65 | 108.9 | 126.9 | 85.8 |
|  | 1014109  (post) | High grade serous carcinoma | T | 3 | 1 | 40 | 172.5 | 128.8 | 134.0 |
| 8 | 1014201  (pre) | High grade serous adenocarcinoma | T |  |  | 35 | 134.8 | 145.3 | 92.7 |
|  | 1014201  (post) | Residual high-grade serous adenocarcinoma | T | 3 | 1 | 45 | 170.6 | 141.6 | 120.5 |
| 9 | 1014478  (pre) | High grade serous adenocarcinoma | T |  |  | 100 | 100.9 | 151.5 | 66.6 |
|  | 1014478 (post) | High grade serous adenocarcinoma | T | 3 | 1 | 65 | 120.5 | 155.3 | 77.6 |
| 10 | 1016196  (pre) | Serous adenocarcinoma | T |  |  | 45 | 138.0 | 137.6 | 100.3 |
|  | 1016196 (post) | High-grade serous papillary carcinoma | T | 3 + Bev | 1 | 50 | 120.2 | 130.9 | 91.8 |
| 11 | 1014249  (pre) | Adenocarcinoma consistent with ovarian serous carcinoma | T |  |  | 35 | 99.9 | 176.2 | 56.7 |
|  | 1014249  (post) | High grade serous carcinoma | T | 6 | 2 | 40 | 90.0 | 112.8 | 79.8 |
| 12 | 1014386  (pre) | High grade serous carcinoma | T |  |  | 75 | 131.1 | 127.0 | 103.3 |
|  | 1014386 (post) | High-grade serous carcinoma | T | 6 | 1 | 70 | 121.1 | 115.4 | 105.0 |
| 13 | 1016266  (pre) | High-grade serous carcinoma | T |  |  | 60 | 72.1 | 120.9 | 59.7 |
|  | 1016266  (post) | High-grade serous adenocarcinoma | T | 6 | 2 | 100 | 123.1 | 139.3 | 88.3 |
| 14 | 3000276  (pre) | Adenocarcinoma consistent with ovarian serous carcinoma | T |  |  | 80 | 106.9 | 149.6 | 71.5 |
|  | 3000276  (post) | High-grade serous adenocarcinoma | T | 6 | 1 | 60 | 110.5 | 141.3 | 78.2 |

Abbreviations used: Pre = initial tumor biopsy; post = tissue removed at cytoreductive surgery following neoadjuvant treatment, as specified; T = Tissue, A = Ascites, or C-T = carboplatin-paclitaxel; Bev = bevacizumab; qualitative PAX8+ (%) = pathologist report; quantification performed using Aperio Image Scope software (Leica Biosystems, version 12.3.0.5056).
