## Supplemental Table S14 for "FAK activity sustains intrinsic and acquired ovarian cancer resistance to platinum chemotherapy"

**Table S9. Primers used for quantitative PCR.**

| Name | Sequence |
| --- | --- |
| musRS17F | ATGACTTCCACACCAACAAGC |
| musRS17R | GCCAACTGTAGGCTGAGTGAC |
| musSusd6F | CATAGTGGCTTCCACTGCCA |
| musSusd6R | CCTGTTCACGCCTGCTATGA |
| musBbc3F | GCGGCGGAGACAAGAAGAG |
| musBbc3R | AAGGGGAGGAGTCCCATGAA |
| musBaxF | GAGCTGCAGAGGATGATTGC |
| musBaxR | CTTGGATCCAGACAAGCAGC |
| musFasF1 | GAGAGTTTAAAGCTGAGGAGGC |
| musFasR1 | TCATCGCAGAGTGTGCATCTT |
| musTigarF | CCTTGACCGTTATCCGCCAT |
| musTigarR | GCATCTACGCCTTGTCCTTG |
| musFuca1F | ACCTCGGCCACAAAGATAACA |
| musFuca1R | TCAGAGTCCACGCAAACTCC |
| musPank1F | GCGCGCAACACGAGC |
| musPank1R | GGGAATGGCGGCCTGTT |
| musFdxrF | TGCTCAGCAATCGAGGAGTC |
| musFdxrR | GAGACTTCCTCAGCATCCAGC |
| musPrkab1F | CACGGGCATCTCTTGTGACC |
| musPrkab1R | TACCGGTGTGTTGCACTGAG |
| musDram1F | CAGCTTCTTGGTCCGACGAG |
| musDram1R | CGTAGCTGCGCCAAGAAATG |
| musRrm2bF | CTGTTCAAATCGAGCAGGAGT |
| musRrm2bR | GCCATGACTGCAAATCGCTG |
| musPolhF | CCTCGCTATGACGCTCACAA |
| musPolhR | AGGAGGGGACCACTCAGTTT |
| musXpcF | ATTCCAGGGATTGCGTGCAT |
| musXpcR | TCCCAAACTCATTCCGAGGC |
| musPcnaF | AAAGATGCCGTCGGGTGAAT |
| musPcnaR | TCTATGGTTACCGCCTCCTCT |
| musGadd45aF | TGGTGACGAACCCACATTCA |
| musGadd45aR | CGGGAGATTAATCACGGGCA |
| musBtg2F | CGCACTGACCGATCATTACAA |
| musBtg2R | GGATCAACCCACAGGGTCAG |
| musSfnF | ACAGGCCGAACGGTATGAAG |
| musSfnR | GTACTCTTTCACCTCGGGGC |
| musCdkn1aF | GCAGACCAGCCTGACAGATTT |
| musCdkn1aR | TGGGCACTTCAGGGTTTTCT |
| musSesn1F | CAGCTGCTGGATCGTAGCTT |
| musSesn1R | ACATGGACCTTCTCAGAGTGC |
| musSesn2F | CACTTCCGCCACTCAGAGAA |
| musSesn2R | CCCCAGCATGTGGTGGATAG |
| musCcng1F | GGCGCTATCTATCCTTGCGT |
| musCcng1R | TCTCGGCCACTTATCTTGGA |
| musPpm1dF | CCACCAATCAAGTCACCGGA |
| musPpm1dR | GCATTACTGCGAACAAGGGC |
| musMdm2F | ACGATGGCGTAAGTGAGCAT |
| musMdm2R | CTGTGACCCGATAGACCTCA |
